# Training-Induced Circuit-Specific Excitatory Synaptogenesis is Required for Effort Control

**DOI:** 10.1101/2021.04.20.440613

**Authors:** Francesco Paolo Ulloa Severino, Oluwadamilola Lawal, Kristina Sakers, Shiyi Wang, Namsoo Kim, Alexander Friedman, Sarah Johnson, Chaichontat Sriworarat, Ryan Hughes, Scott Soderling, Il Hwan Kim, Henry Yin, Cagla Eroglu

**Author notes:** Corresponding authors e-mail addresses.

## Abstract

Synaptogenesis is essential for circuit development; however, it is unknown whether it is critical for the establishment and performance of goal-directed voluntary behaviors. Here, we show that operant-conditioning via lever-press for food reward-training in mice induces excitatory synapse formation onto a subset of Anterior Cingulate Cortex neurons projecting to the dorsomedial striatum (ACC_->DMS_). Training-induced synaptogenesis is controlled by the Gabapentin/Thrombospondin receptor α2δ-1, which is an essential neuronal protein for proper intracortical excitatory synaptogenesis. Using germline and conditional knockout mice, we found that deletion of α2δ-1 in the adult ACC_->DMS_ circuit diminishes training-induced excitatory synaptogenesis. Surprisingly, this manipulation did not impact learning but instead resulted in a profound increase in effort exertion without affecting sensitivity to reward value or changing contingencies. Bidirectional optogenetic manipulation of ACC_->DMS_ neurons rescued or phenocopied the behaviors of the α2δ-1 cKO mice highlighting the importance of synaptogenesis within this cortico-striatal circuit in regulating effort exertion.

## Introduction

Goal-directed behaviors are executed to obtain desirable outcomes, such as food rewards. These complex voluntary behaviors establish when motivated individuals learn to associate a set of actions with its desirable outcome^1^. An important aspect of goal-directed behaviors is effort, which can be described as the motor and cognitive resources allocated to action performance. Learned behaviors need to be adaptable to changing action-outcome contingencies, because often the effort required to reach the desired outcome changes. Maladaptive behaviors such as spending too much effort on ineffective and demanding actions can be detrimental to survival^2, 3^.

Synapses are the smallest units of neuronal circuits that control behaviors. The molecular mechanisms regulating synaptogenesis are highly complex. Importantly, mutations in genes encoding for synaptic or synaptogenic proteins are linked to a large number of brain diseases^4^ with severe cognitive decline, sensorimotor and memory deficits^5–12^. Majority of the synaptic structures are established during development; however, synaptic connectivity is not stagnant and is dynamically modified throughout the life span^13–15^. This experience-dependent remodeling of synapses is thought to underlie cognitive processes such as learning and memory^16–21^.

Long-term synaptic plasticity is widely accepted as the mechanism underlying synaptic remodeling associated with learning^21–24^. However, whether synaptogenesis is involved in the establishment and performance of complex behaviors is unclear. In this study, in freely moving animals, we investigated whether new synapse formation occurs during training and if it is required for learning reward-based behaviors. To do so, we used an instrumental operant-conditioning task in mice^25, 26^. In this behavioral paradigm, a mouse first learns to perform a specific action (i.e., press a lever) to earn a reward (i.e., food pellet). Once the mouse associates the action and outcome, the contingency can be changed either by increasing the number of lever presses required to receive the reward or by manipulating the relationship between lever press and reward. The mice adapt their rate of lever pressing based on these changing contingencies. For example, mice increase their effort, when the number of presses required to achieve a reward is increased, but they would diminish lever press rate if the task becomes too demanding ^27, 28^.

Previous research, utilizing similar operant-conditioning paradigms, identified brain regions within the basal ganglia circuits to be responsible for controlling instrumental actions, and revealed how disruption of these circuits affects learning and performance^29–31^. For example, learning the action-outcome relationship requires the dorsomedial striatum (DMS)^29, 32^, which is a hub for many cortical inputs^33, 34^. Particularly, the inputs coming from the prefrontal cortex (PFC) to the DMS play critical roles in instrumental training, such as learning the action- outcome contingency^31, 35, 36^, evaluation of an outcome’s value^30, 37, 38^, and deciding between different actions or outcomes^39, 40^. However, the cellular and molecular mechanisms underlying the remodeling of these cortico-striatal circuits during learning and how their functions control goal-directed actions are unknown. Closing these fundamental knowledge gaps is needed to determine druggable molecular targets to treat brain disorders, such as obsessive compulsive disorder (OCD), autism spectrum disorder (ASD) and Alzheimer’s disease (AD), in which goal-directed behaviors are impaired^41–44^.

Which cortical regions control learning and establishment of instrumental behaviors? Does training induce cortical synaptogenesis? Are the newly formed synapses required for establishment of learned goal-directed actions? Here, we addressed these questions by applying molecular and anatomical tools, circuit-specific gene modifications, optogenetics and quantitative behavioral techniques. By investigating the immediate early gene c-Fos expression as a marker for structural synaptic plasticity and neuronal activity^45–48^, we found the Anterior Cingulate Cortex (ACC) to be significantly activated by the instrumental training. This knowledge allowed us to identify an ACC_->DMS_ connection that is controlling instrumental behavior performance.

In humans PFC, a brain region containing ACC, is associated with cognitive control, the ability to flexibly adjust our behaviors and allocate effort towards our goals^49–52^. In rodents, the ACC has been suggested to regulate learning^53^, reward and action monitoring^54^, decision making^55^, impulsivity^56^ and selection of behavioral strategy^57^, because the neuronal activity in the ACC correlates with these aspects of complex behaviors. However, the cellular and molecular mechanisms utilized by the ACC circuits to achieve such roles are unknown. To determine if new synapse formation within the ACC_->DMS_ circuit we identified is critical for the establishment and performance of goal directed actions, we took a genetic approach to block synaptogenesis specifically in these neurons. To do so, we targeted the Gabapentin receptor α2δ-1 is a type-I membrane protein encoded by Cacna2d-1^58^.

α2δ-1 is essential for the proper formation and maturation of intracortical excitatory synapses during development^59^. α2δ-1 (Cacna2d-1) is also highly expressed in the adult mouse cortex^60^. It was first identified as a subunit of voltage-gated calcium channels^61^. Then it was shown to interact with several synaptic and synaptogenic proteins^58, 62^, including the astrocyte-secreted Thrombospondins (TSPs)^59, 63^. The synaptogenic function of α2δ-1 is mediated by Rac-1- signaling within the dendrites of cortical neurons^59^ and it is independent of its role in calcium channel trafficking^63–65^. Despite its known critical functions in the development and maturation of synapses, whether α2δ-1 controls adult synaptogenesis and contributes to learning and performance of complex behaviors remain poorly understood. Our findings identify α2δ-1- dependent adult synaptogenesis as a critical cellular mechanism controlling the adaptability of voluntary goal-directed actions. Surprisingly, training-induced synapse formation is not required for learning but for the control of effort exertion through action sequence modulation.

## Results

### Lever press task induces the establishment of lever press bouts and increases immediate early gene expression in the ACC

To model operant behaviors in mice, we used an instrumental conditioning task, in which mice learn to press a lever for a food reward. In this paradigm, mice first learn the relationship between an action (lever press, LP) and the desired outcome (food reward). Following this initial phase, the animal’s performance can be improved by increasing the number of LPs required for each reward (“Fixed Ratio” Fig. 1a).

**Figure 1.**
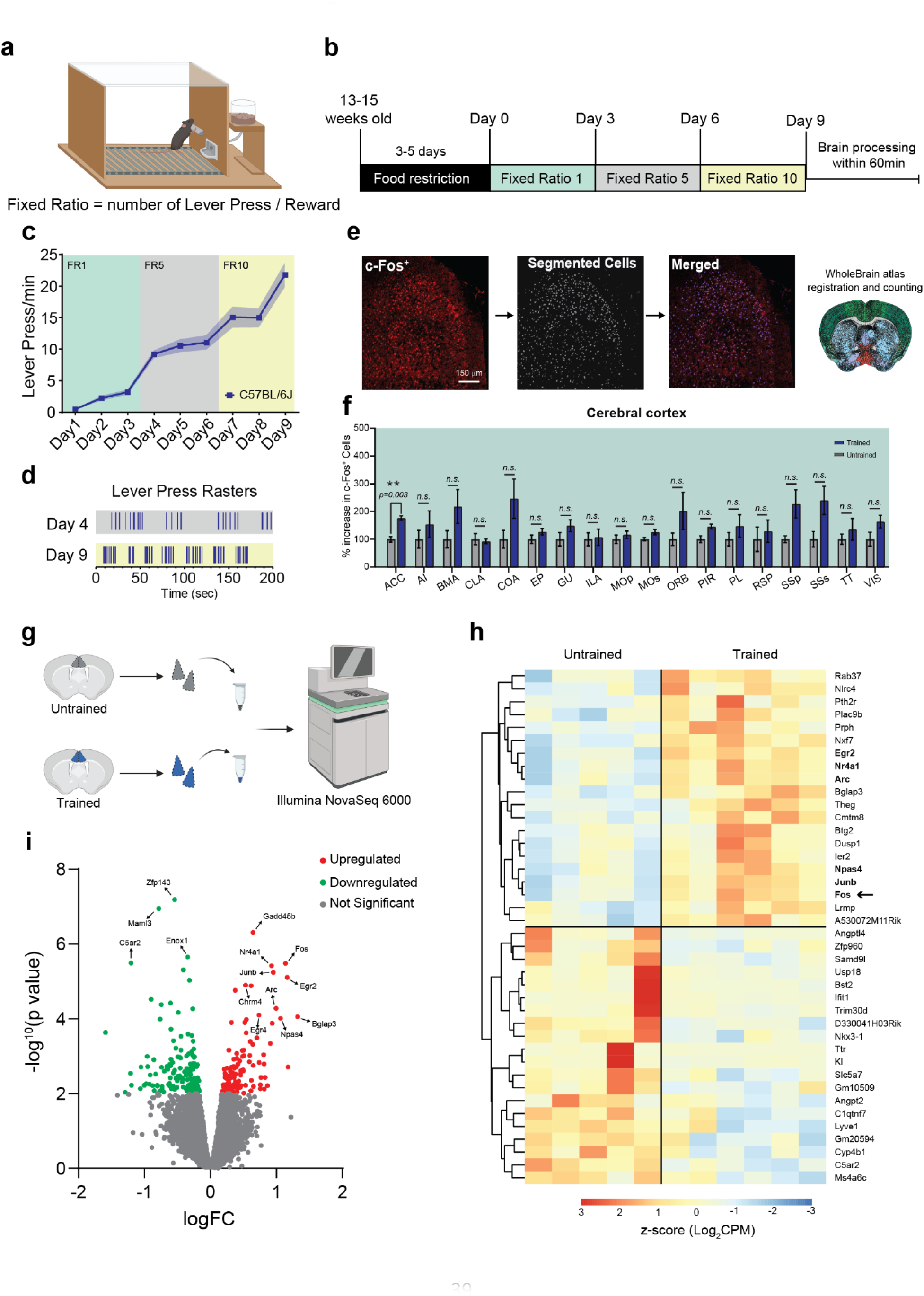
Action sequence engages the Anterior Cingulate Cortex during instrumental actions. **a**, Representation of the Skinner box used for training and testing mice. **b**, Schematic of the schedules used for the operant training and testing. **c**, Lever press (LP) rate for trained C57BL/6J mice (n=17 mice, 8 males, and 9 females) across the 9 days of training (Day 1 = 3 ± 0.4 LP/min, Day 9 = 22 ± 1.9 LP/min; *One-way ANOVA for repeated measures,* main effects of *Days* [F (8, 128) = 74.86, p < 0.0001] and *Subject* [F (2.13, 17.04) = 10.30, p = 0.001]. **d**, Representative lever press raster plots for FR5 day 4 and FR10 day 9 of LP performance. **e**, Flow chart and example images of the c-Fos^+^ cell segmentation and quantification. **f**, Bar plot of c-Fos^+^ cells for the cerebral cortex regions. *Multiple unpaired t-test with Welch correction. Multiple comparisons using Holm-Sidak method; alpha = 0.05 for adjusted p-value*. ACC (176.203 ± 8.26%, [t (9.684) = 5.906]). n = 4-6 mice per condition. **g**, Schematic representation of the ACC microdissection and RNA-seq from trained and untrained mice. **h**, Heat map of the top 20 differentially expressed genes (DEG) in the Trained and Untrained groups that are ranked based on the z-score of the log_2_CPM. Bolded genes are the IEGs overexpressed in Trained mice. **i**, Volcano plot for DEG is organized based on their logarithmic fold change (logFC) and p-value (-log_10_p value). Arrows signify genes of interest.

We trained male and female wild-type (WT, C57BL/6J) mice using a fixed ratio (FR) schedule to identify brain regions responsible for the learning and performance of the LP task. Mice were first trained on an FR1 schedule (1 lever press (LP)/1 reward, days 1-3), then on an FR5 (5 lever presses/1 reward, days 4-6), and finally on an FR10 schedule (10 lever presses/1 reward, days 7-9) (Fig. 1b, see Methods for details). An untrained control group (age- and sex-matched) was also food-restricted and housed in the same training chamber for an equivalent number of days and durations as the trained mice. However, the untrained mice were given free access to the same amount of food rewards without pressing the lever.

Throughout the training, the mice learned to press the lever efficiently, which is reflected by the significant increase in the number of LPs/min between the first day of training (Day 1-FR1) compared to the last day (Day 9-FR10) (Fig. 1c). Apart from the increased number of LPs/min, we utilized the probability distribution of inter-press intervals (IPI, Supplementary Fig. 1a) to identify and analyze the lever press bouts across days of training as a parameter of improved task performance (Fig. 1d and Supplementary Fig. 1b). Quantification of bout duration, number of LPs per bout, mean inter-press interval (IPI) within each bout and inter-bout interval (IBI) from day 4 through day 9 show an overall improvement of performance over time (Supplementary Fig. 1c-f). We did not find any sex differences in the performance of this task (Supplementary Fig. 1g). These data show that in this paradigm mice of either sex learn the action-outcome contingency and efficiently adapt to contingency changes (i.e., transitioning to FR5 and FR10) by performing bouts of discrete number of LP.

Which brain regions are critical for learning and performance of the lever press task? To answer this question, we first analyzed the immediate early gene (IEG) c-Fos expression, as a cellular marker of increased neuronal activity, in several brain regions ^46, 66^ after the FR10 schedule. To capture the c-Fos protein at its highest levels^46^, we processed the brains of the trained and untrained mice within 60 minutes after completing the last session on day 9 (Fig. 1b). Coronal brain sections from trained and untrained mice corresponding to 4 forebrain Bregma coordinates (Supplementary Fig. 1h, posterior-anterior: -0.3; +0.4; +1.7; +2.2) were stained for c-Fos and DAPI (nuclear DNA marker). The c-Fos^+^/DAPI^+^ cells (hereafter named c-Fos^+^) were imaged from the entire coronal brain sections and segmented using U-Net machine-learning algorithm (https://github.com/ErogluLab/CellCounts, see Methods for details). With the Whole Brain Software ^67^, we mapped the segmented c-Fos^+^ cells onto the Allen brain atlas for the corresponding Bregma coordinates (Fig. 1e and Supplementary Fig. 1i). This analysis revealed a significant increase in the total number of c-Fos^+^ cells in trained compared to untrained mice (Supplementary Fig. 1j). The 42 detected brain regions were grouped based on their classification as regions belonging to the cerebral cortex (Fig. 1f), cerebral nuclei (Supplementary Fig. 1k), or interbrain (Supplementary Fig. 1l). A significant increase in c-Fos^+^ cells after training was detected in the Anterior Cingulate Cortex (ACC) and the Magnocellular Nucleus (MA). Previous work has implicated the prefrontal cortex in the learning and performance of instrumental actions, decision-making in cost-benefit tasks, and behavioral flexibility ^31, 39, 40, 57, 68–70^. We, therefore, focused on the ACC for the remainder of our study because our findings suggested that the ACC, among the prefrontal cortex regions, is strongly and selectively activated by instrumental training.

To further investigate and confirm the changes in gene expression due to the training, we micro- dissected the ACC from trained and untrained mice and performed RNA-sequencing (RNA-seq) (Fig. 1g). Of the seventy-five differentially expressed genes (DEGs) identified, thirty-two were significantly upregulated in the ACC of trained mice, and 43 were downregulated (i.e., 1.5-fold change compared to untrained animals, at a nominal p-value of p <0.01, Fig. 1h,i).

We found several immediate early genes (IEGs) among the genes upregulated by training, such as *Fos, Jun, Npas4, Arc, Nr4a1*, *Egr2*, and *Egr4* (Fig. 1h,i), known for their role in synaptic plasticity and memory-related processes ^71–73^. Indeed, GO term analyses on DEGs showed a significant (FDR ≤ 0.05) enrichment ratio (ER) for genes involved in cognitive processes (ER = 12.8) and DNA-binding transcription activators (ER = 9.0). These results reveal increased IEG expression in the ACC of trained mice, providing evidence for heightened neuronal activity in this region during the performance of the LP task. Furthermore, because IEGs are involved in the molecular mechanisms underlying long-term structural and functional changes in synaptic circuits during learning and memory ^71, 73, 74^, these findings suggest that LP training induces long-term circuit remodeling in the ACC.

### Excitatory synapses increase in the ACC after training

Closer inspection of cFos^+^ cells in the ACC revealed that instrumental training significantly increases the number of c-Fos^+^ cells in layers 2/3 of the ventral ACC (vACC, Fig. 2a, b). The ACC layers 2/3 and 5 contain neurons that project to the DMS, a striatal region that controls instrumental actions and action sequences^33, 75, 76^. In contrast, there were no significant differences in the numbers of c-Fos^+^ cells between trained and untrained mice in the dorsal ACC (Fig. 2a, b) and the neighboring secondary Motor Cortex (MOs), which also sends projections to the DMS (Supplementary Fig. 2a). These results show that there is a significant increase in the numbers of c-Fos^+^ cells in the L2/3 of the vACC of trained mice and implicate the ACC neurons that project to the DMS as neurons that have increased activity after training.

**Figure 2.**
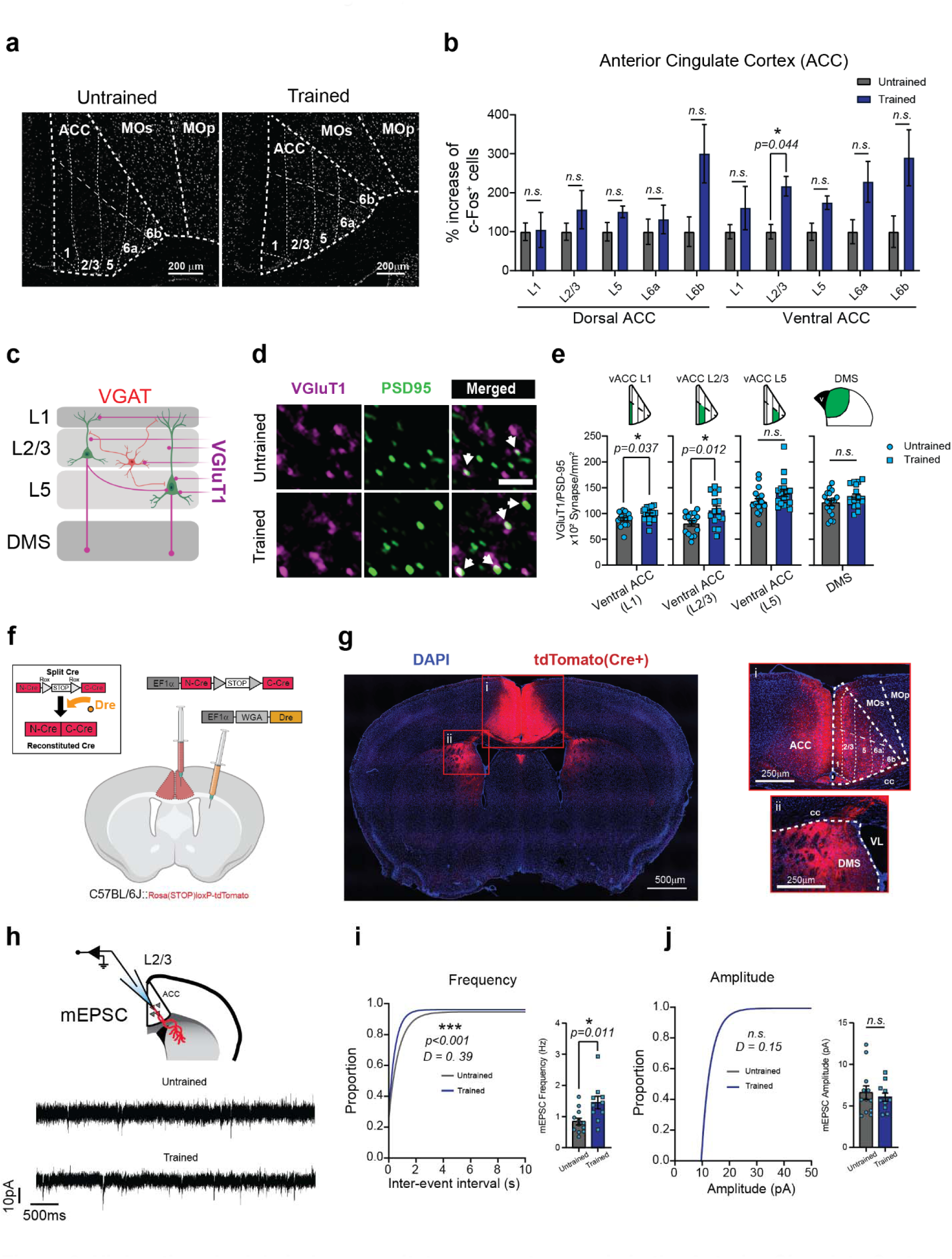
High ratio schedule induces excitatory synaptogenesis in the Anterior Cingulate Cortex. **a**, Example images of the segmented c-Fos^+^ cells across the layers of the ACC. **b**, Layer specific quantification of the c-Fos^+^ cells in the dorsal and ventral ACC in trained and untrained C57BL/6J mice (n = 6 mice per condition (3 males and 3 females); 4 sections per mouse*). Multiple unpaired t-test with Welch correction. Multiple comparisons using Holm-Sidak method; adjusted p-value*. vACC 2/3 [t (9.18) = 3.72]. **c**, Representation of the specific regions in which synaptic analysis was performed for VGlut1 and VGAT synapses. L1, cortical layer 1; L2/3, cortical layers 2/3’ L5, cortical layer 5; DMS, dorsomedial striatum. **d**, Representative images from Untrained and Trained mice that were stained with VGluT1 and PSD95 antibodies. The *arrows* in the merged channel indicate co-localized puncta. *Scale bar 2 µm*. **e**, Quantification of VGluT1/PSD95 co-localized puncta density in the ACC and DMS. *n = 4 mice per condition, 3 images per mouse. Unpaired Two-tailed t-test.* L1 Untrained (89.1 ± 3.02), Trained (98.3 ± 2.97) [t (34) = 2.2]; L2/3 Untrained (81.0 ± 5.34), Trained (107 ± 7.92) [t (28) = 2.7]; L5 Untrained (124 ± 5.58), Trained (140 ± 6.87) [t (34) = 1.9]. **f**, Schematic representation of the injections for both EF1a- WGA-Dre and EF1a-N-Cre-*rox-STOP-rox-*C-Cre viruses. **g**, Tile scan image of a coronal brain section from a C57BL/6J tdTomato^+^ mouse, showing labeled ACC_->DMS_ neurons. Magnification of the ACC (i) and DMS (ii) of a mouse injected with the WGA-Dre and the Rox-Cre viruses. **h**, Schematic representation of mEPSC recording from ACC_->DMS_ neurons and representative traces from Untrained and Trained mice. **i**, Comparison between Untrained (n = 12 cells from 4mice) and Trained (n = 11 cells from 4 mice) WT mice of the cumulative distribution of inter-event interval, *Kolmogorov-Smirnov test*; average mEPSC frequency, Untrained (0.84 ± 0.11) and Trained (1.45 ± 0.20). *Unpaired Two-tailed t-test [t (20) = 2.78]*. **j**, Cumulative distribution of amplitude *Kolmogorov-Smirnov test (p = 0.474);* average mEPSC amplitude, Untrained (6.59 ± 0.81) and Trained (6.05 ± 0.49). *Unpaired Two-tailed t-test [t (21) = 0.56].* For all graphs: Data shown as mean ± s.e.m. alpha = 0.05.

We postulated that training promotes a net increase in excitatory synapses onto the neurons in this region. To test this hypothesis, we compared the numbers of excitatory or inhibitory synaptic structures in the vACC L2/3 and L5 and the synaptic zone (layer 1, L1) of the vACC from trained and untrained mice. The synaptic zone in L1 harbors the apical dendrites from L2/3 and L5 neurons, which receive many synapses. In addition, we analyzed synapse numbers in the DMS, the axonal target for ACC neurons (Fig. 2c). To visualize and quantify structural synapses, we used an established protocol^77^ that marks synapses as the juxta-positioning of pre and postsynaptic markers. This method takes advantage of the close proximity of pre and postsynaptic markers and of the resolution limit of light microscopy. The pre and postsynaptic proteins are in distinct neuronal compartments (axons and dendrites, respectively); however, due to their close proximity at synapses they appear as partially co-localized.

We used the Vesicular Glutamate Transporter 1 and the PostSynaptic Density protein 95 (VGluT1/PSD95) or the Vesicular GABA Transporter and gephyrin (VGAT/Gephyrin) to mark the respective pre and postsynaptic compartments of excitatory or inhibitory synapses (Fig. 2d and Supplementary Fig. 2d). We found an increase in the VGluT1/PSD95-positive excitatory synapse density in the L1 and L2/3 of the vACC of trained mice compared to untrained controls (Fig. 2e). We excluded that the observed increase was due to an enlargement of the pre- and post-synaptic side as we found no changes in the puncta size after training (Supplementary Fig. 2c). Moreover, no significant changes in synapse densities were observed in the vACC L5 or the DMS (Fig. 2e), or within the cortical region adjacent to ACC, the MO (Supplementary Fig. 2b). Finally, training did not alter the numbers of VGAT/Gephyrin-positive inhibitory synapses in any of these regions (Supplementary Fig. 2e). These data show that the training and improved performance of instrumental actions is accompanied by a significant increase in the numbers of VGluT1/PSD95-positive excitatory synapses in the L2/3 of the vACC.

Next we tested whether the increase in the numbers of structural excitatory synapses we observed reflects a net functional increase in excitatory inputs onto the ACC neurons that are projecting to the DMS (here after called, ACC_->DMS_ neurons). To do so, we recorded miniature excitatory postsynaptic current (mEPSC) from ACC_->DMS_ neurons of the trained and untrained mice. To label ACC_->DMS_ neurons, we used a viral approach that relies on the combined functions of two adeno-associated viruses (AAVs) ^78^. The first AAV is injected bilaterally into the DMS and expresses the Dre-recombinase protein conjugated to Wheat Germ Agglutinin (WGA). WGA is a lectin that retrogradely transports across synapses^79^. Hence, WGA-conjugated Dre recombinase is expressed by DMS neurons and transferred to all the neurons that send synapse onto the DMS. A second AAV is bilaterally injected into the ACC and contains the Cre- recombinase coding sequence; however, Cre protein expression is interrupted by a Rox-flanked STOP cassette (N-Cre-*rox-STOP-rox*-C-Cre, Fig. 2f). This strategy ensures that Cre is only expressed in the ACC_->DMS_ neurons, because the retrogradely transported WGA-Dre allows the recombination of the rox-STOP-rox codon within the Cre recombinase (Fig. 2f). We verified the efficiency and specificity of this approach to target ACC_->DMS_ neurons using a Cre-reporter mouse line, Rosa (STOP)loxP-tdTomato^80^ (Fig. 2g).

This approach allowed us to perform mEPSC recordings specifically in the L2/3 ACC_->DMS_ neurons (Fig. 2h) in trained and untrained mice. In line with an increase in the number of excitatory synapses, we found a functionally significant increase in the frequency, but not in the amplitude, of mEPSCs in the L2/3 ACC_->DMS_ neurons of the trained mice compared to untrained controls (Fig. 2i, j). These findings strongly suggest that operant training induces excitatory synaptogenesis onto ACC_->DMS_ neurons. These findings also implicate excitatory synapse formation as a critical step which is necessary for the learning and performance of instrumental actions.

### Training-induced excitatory synapse formation in the ACC requires the synaptogenic neuronal receptor **α**2**δ**-1

Next, we took a genetic approach to determine if training-induced synapse formation is required for instrumental learning and performance. *Cacna2d1* encodes for the synaptogenic neuronal receptor α2δ-1, which is required for intracortical synaptogenesis during development in the sensory cortical regions ^59^. By mining single-cell RNA sequencing data from adult mouse brain ^81^ we found that, out of all the genes, *Cacna2d1* is also highly expressed in the prefrontal cortex (Supplementary Fig. 3a) with the pyramidal neurons in the L2/3 having the highest expression levels (Supplementary Fig. 3b). To investigate whether training-induced VGluT1/PSD95-positive synapse formation is dependent on α2δ-1, we bred α2δ-1 heterozygous mice and analyzed the numbers of synapses in the vACC of adult trained and untrained α2δ-1 WT and KO offspring (Fig. 3a).

**Figure 3.**
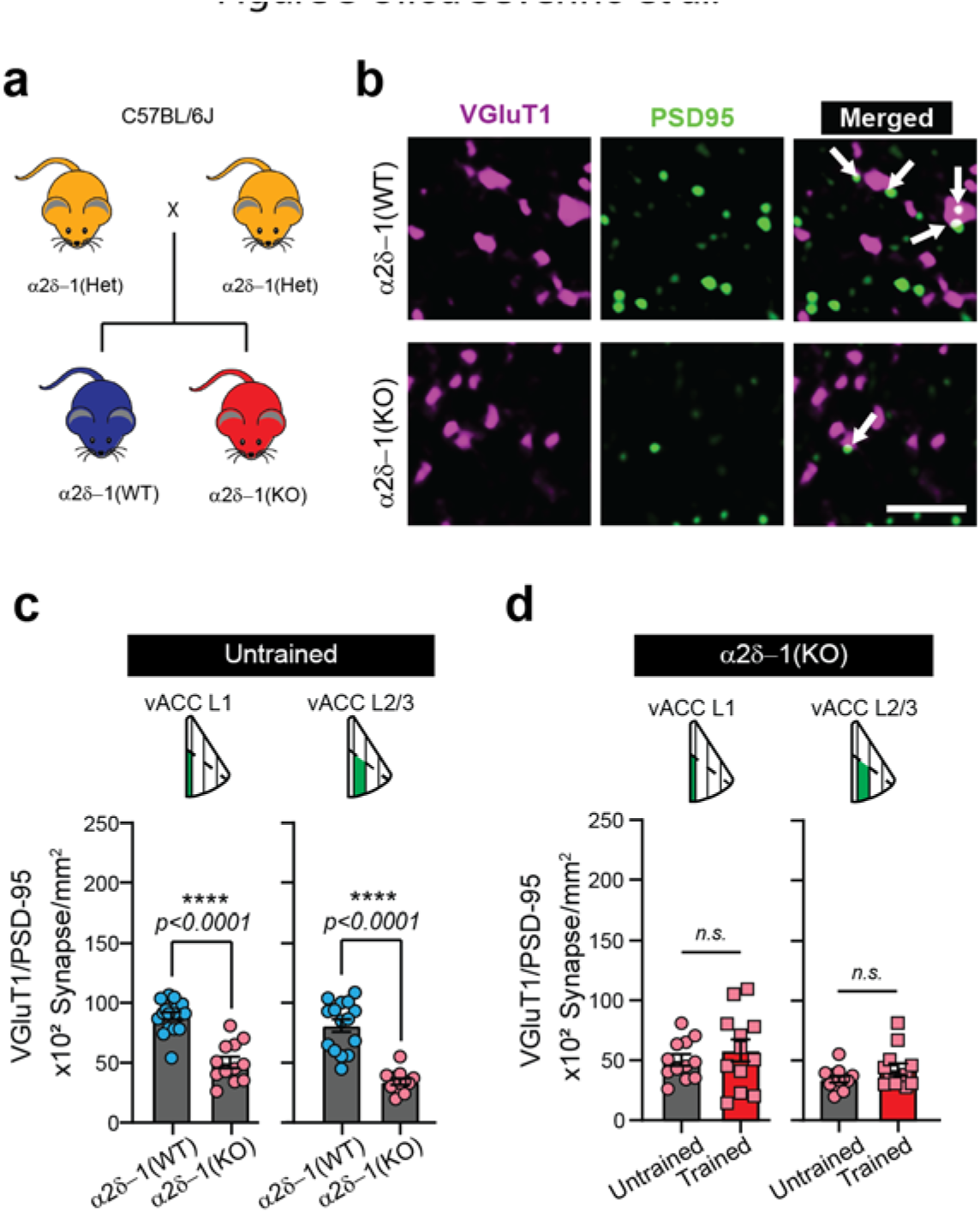
VGlut1-PSD95 synapse formation in the Anterior Cingulate Cortex is regulated by the Thrombospondin receptor α2δ-1. **a**, Breeding scheme for the generation of α2δ-1 WT and KO mice. **b**, Representative images of VGluT1/PSD95 staining in the ACC of α2δ-1 WT and KO mice. The *arrows* in the merged channel indicate co-localized puncta. Scale bar 2µm. **c**, Comparison between untrained α2δ-1 WT and KO mice in L1 (WT 89 ± 3.0; KO 50 ± 4.8), Unpaired Two-tailed t-test [t (28) = 7.2] and L2/3 (WT 81 ± 2.7; KO 34 ± 2.7), Unpaired Two-tailed t-test [t (25) = 7.3]. *n = 4-6 mice per condition and genotype (n = 3 images per mouse).* **d**, Layer-specific comparison of untrained and trained α2δ-1 KO in the ventral ACC. L1 (Untrained 50 ± 4.8; Trained 58 ± 9.1), Unpaired *Two-tailed t-test [t (22) = 0.77]*. L2/3 (Untrained 34 ± 2.7; Trained 43 ± 4.8), Unpaired *Two-tailed t-test [t (22) = 1.5].* For all graphs: *Multiple unpaired t-test with Welch correction. Multiple comparisons using Holm-Sidak method; alpha = 0.05 for adjusted p-value*. Data shown as mean ± s.e.m.

We found that loss (KO) of α2δ-1 already severely reduces the synapse density in the vACC of the untrained mice (∼50% reduction, Fig. 3b, c). Next, we tested if loss of α2δ-1 affects training-induced synaptogenesis. Here, we compared the numbers of VGluT1/PSD95-positive synapses among untrained and trained α2δ-1 KO mice. The training-induced increase in VGluT1/PSD95 positive synapses in L1 and L2/3 were abolished in α2δ-1 KO mice (Fig. 3d). The average size of the pre-synaptic marker VGlut1 were unaltered (Supplementary Fig. 3c) whereas post- synaptic marker PSD95 puncta size were decreased in α2δ-1 KOs (Supplementary Fig. 3d), a finding in line with the important role of α2δ-1 in spinogenesis^59^. Our findings show that diminished α2δ-1-signaling reduces excitatory synapse numbers in the vACC of adult mice and prevents training-induced excitatory synaptogenesis in the L1 and L2/3 of the vACC. These results suggest that α2δ-1 KOs can be utilized as a genetic tool to test the role of cortical synaptogenesis on instrumental learning and performance.

### Loss of **α**2**δ**-1-signaling does not impair learning of instrumental actions but causes an increase in effort exertion

We examined whether α2δ-1 KO mice would exhibit deficits in the LP task (Fig. 4a). Surprisingly, there were no genotype or sex differences in the ability of the KO mice to learn and perform the FR lever press schedules (Fig. 4b and Supplementary Fig. 4a, b). Analysis of the lever press patterns established during training confirmed a significant improvement in performance across days in both genotypes (Fig. 4b and Supplementary Fig. 4c-f). These findings suggest that training-dependent, α2δ-1-mediated excitatory synaptogenesis is not necessary to establish instrumental actions.

**Figure 4.**
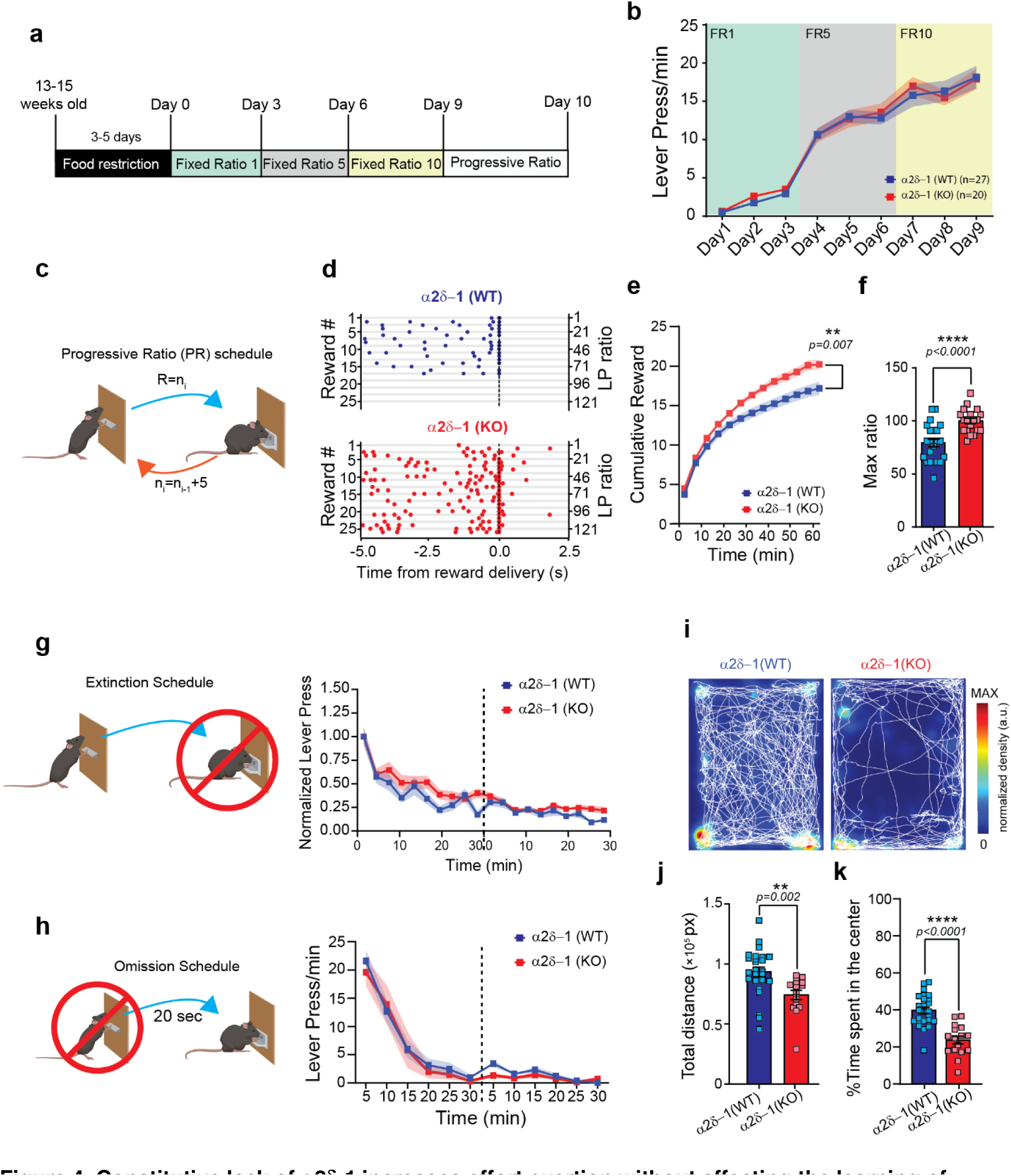
Constitutive lack of α2δ-1 increases effort exertion without affecting the learning of instrumental actions. **a**, Schematic representation of the training schedule used for the α2δ-1 WT and KO mice. **b**, Lever press (LP) rate for the 9 days on the FR schedule for α2δ-1 WT (n = 27 mice; 14 male and 13 female) and KO (n = 20 mice; 10 male and 10 female). *RM Two-way ANOVA. Main effects of Days [F (2.536, 114.1) = 196.4, p < 0.0001], no effect of Genotype [F (1, 45) = 0.04814, p = 0.827] nor interaction [F (8, 360) = 0.4819, p = 0.869].* **c**, Schematic representation of the Progressive Ratio (PR) schedule. The value (n) of the ratio (R) increment of 5 for every received reward (i), starting with R=1. **d**, Representative peri-reward raster histograms of LP for α2δ-1 WT and KO mice. **e**, Cumulative reward count over the PR session (bin=5 min) for α2δ-1 WT (n = 22; 17.2 ± 0.8 rewards) and KO (n = 19; 20.2 ± 0.6 rewards) animals. *Two-way ANOVA for repeated measures. Main effects of Genotype [F (1, 39) = 8.060] and Time [F (1.683, 65.65) = 636.6, p < 0.0001] and interaction [F (12, 468) = 7.220, p < 0.0001].* **f**, Max Ratio for α2δ-1 WT (n = 22; Max Ratio = 79.1 ± 3.8 and KO (n = 19; Max Ratio = 99.9 ± 2.6). *Unpaired Two-tailed t-test [t (39) = 4.4].* **g**, Schematic representation of the extinction schedule. α2δ-1 WT (n = 22) and KO (n = 19). The normalized number of lever press is reported for the 2 days of testing (dashed line). *RM Two-way ANOVA. Main effect of time [F (8.158, 318.1) = 31.66, p < 0.0001], no effect of genotype [F (1, 39) = 4.077, p = 0.050] and no interaction [F (19, 741) = 0.9839, p = 0.478].* **h**, Schematic representation of the omission schedule. α2δ-1 WT (n = 17) and KO (n = 8). The lever press/min are reported for the 2 days of testing (dashed line). *RM Two-way ANOVA. Main effect of time [F (3.755, 86.36) = 47.89, p < 0.0001], no effect of genotype [F (1, 23) = 0.4091, p = 0.528] and no interaction [F (11, 253) = 0.2691, p = 0.991].* **i**, Examples of open field patterns of locomotion by α2δ-1 WT and KO mice. **j**, Bar graph of the total distance travelled in pixels for α2δ-1 WT (n = 25; 9.3*10^4^ ± 4.0*10^3^ pixels) and KO (n = 16; 7.4*10^4^ ± 3.8*10^3^ pixels) mice during the open field test. *Unpaired Two-tailed t-test [t (39) = 3.2].* **k**, Percent time spent in the center of the arena by α2δ-1 WT (n = 25; 40 ± 1.7%) and KO (n = 16; 24 ± 2.1%) mice. *Unpaired Two-tailed t-test [t (39) = 5.9].* For all graphs: Data shown as mean ± s.e.m. alpha = 0.05.

What then is the role of synaptogenesis in the ACC? To answer this question, we next tested the performance of WT and KO mice in a Progressive Ratio (PR) paradigm. This schedule evaluates the motivational state of animals by adjusting the effort requirement for a reward^82, 83^, and it is the standard paradigm to study effort in both humans and rodents^37, 84–86^ eliminating the confounding effects of choice-based tasks. During PR test, the number of lever presses required to receive one food reward is progressively increased by an increment of 5 (i.e., 1 LP for the first reward, 6 LPs for the second, 11 LPs for the third, and so on…, Fig. 4c). Mice are expected to stop pressing or reduce their press rates greatly when the number of presses required to achieve the reward becomes too high.

We found that in the PR schedule as the task became difficult, there was a clear difference between how WT and α2δ-1 KO mice responded to the higher LP demand. Surprisingly, throughout the PR session, the performance of α2δ-1 KO mice was higher than the WT control (Fig. 4d). As a result, the KO mice received more rewards and reached a significantly higher maximum ratio by the end of the PR session (Fig. 4e, f). Bout analysis showed that the bout properties remain unaltered in α2δ-1 KO mice, the difference was found in the Inter-Bout interval (IBI) which was higher in α2δ-1 KO mice (Supplementary Fig. 4g-j). These results indicate that in α2δ-1 KO mice cannot effectively control effort exertion during a demanding task due to more frequently repeated bouts.

However, to reach this conclusion several other possible interpretations had to be ruled out, such as the establishment of persistent, impulsive, or hyperactive behaviors in the α2δ-1 KO mice. To do so, both α2δ-1 WT and KO mice were tested for their ability to extinguish the LP behavior (Fig. 4g) or to learn the opposite contingency, i.e., “not pressing the lever to get the reward” (Fig. 4h). In both cases, α2δ-1 WT and KO were able to reduce the number of presses, excluding the possibility of action persistence. We also tested the possibility that in α2δ-1 KOs’ sensitivity to reward value could have a role in the alteration of the effort/reward relation. To test this possibility, we performed a devaluation test. Mice were trained as usual up to FR10. On the 10^th^ day of testing, trained mice were pre-fed either with standard chow (valued state) or reward pellets (devalued state of reward). These mice were then tested on a 10min extinction schedule (Supplementary Fig. 4k). WT mice which were in the valued state pressed more compared to when they were in the devalued state. Comparison of α2δ-1 WT and KO mice in a valued versus devalued reward state showed that α2δ-1 KO mice respond similarly to WT mice (Supplementary Fig. 4l). These results show that in α2δ-1 KO mice effort control is impaired; however, these KO mice still are able to extinguish the LP behavior, learn to reverse or diminish their behavior when the reward contingencies are altered.

Finally, to determine if α2δ-1 KO mice had a hyperactive phenotype, which may explain enhanced LP behavior, we tested them in an open field to quantify their overall motor activity in a novel environment (Fig. 4i). On the contrary to hyperactivity, compared to α2δ-1 WT, the α2δ-1 KOs were significantly less active, as shown by the reduced total distance traveled (Fig. 4j). Moreover, α2δ-1 WT mice spent on average 39.8% of their time in the center zone of the open field arena, whereas the α2δ-1 KO mice spent significantly less time in the center (23.7% Fig. 4k). These results reveal that loss of α2δ-1 inhibits the training-induced excitatory synapse formation in the mouse ACC, but it does not affect the learning of instrumental actions. Instead, the α2δ-1 KO mice display a profound increase in effort exertion, suggesting that excitatory synaptogenesis in the adult cortex is involved in effort regulation.

### Conditional deletion of **α**2**δ**-1 from ACC_->DMS_ neurons is sufficient to increase effort exertion

To determine the specific roles of α2δ-1-signaling in the adult ACC_->DMS_ neurons, we utilized α2δ-1(f/f) mice carrying the Cre-reporter Rosa(STOP)loxP-tdTomato (Fig. 5a) and conditionally deleted α2δ-1 selectively from these ACC neurons by utilizing the viral approach described before (Fig. 2f). We trained virally transduced α2δ-1(+/+) or α2δ-1(f/f) mice using the same behavioral paradigm (Fig. 4a). Similar to α2δ-1 global KOs, we found that circuit-specific deletion of α2δ-1 in adulthood does not affect the learning of the lever press task (FR1 schedule in Fig. 5b). During both FR5 and FR10 schedules, we observed a trending, but not significant, increase in the LP rate for the α2δ-1(f/f) compared to the α2δ-1(+/+) mice (Fig. 5b). Analysis of the lever press bouts (Supplementary Fig. 5a-d) showed an overall improvement of both groups across days. However, a significant difference between α2δ-1(+/+) and α2δ-1(f/f) was observed for the number of presses per bout and bout IPI (Supplementary Fig. 5b, c), showing that mice lacking α2δ-1 in ACC_->DMS_ neurons have an alteration of the lever press sequence, with an increased number of presses within bouts. These results suggest a specific function of α2δ-1 in the ACC_->DMS_ neurons in controlling the properties of the lever press bouts.

**Figure 5.**
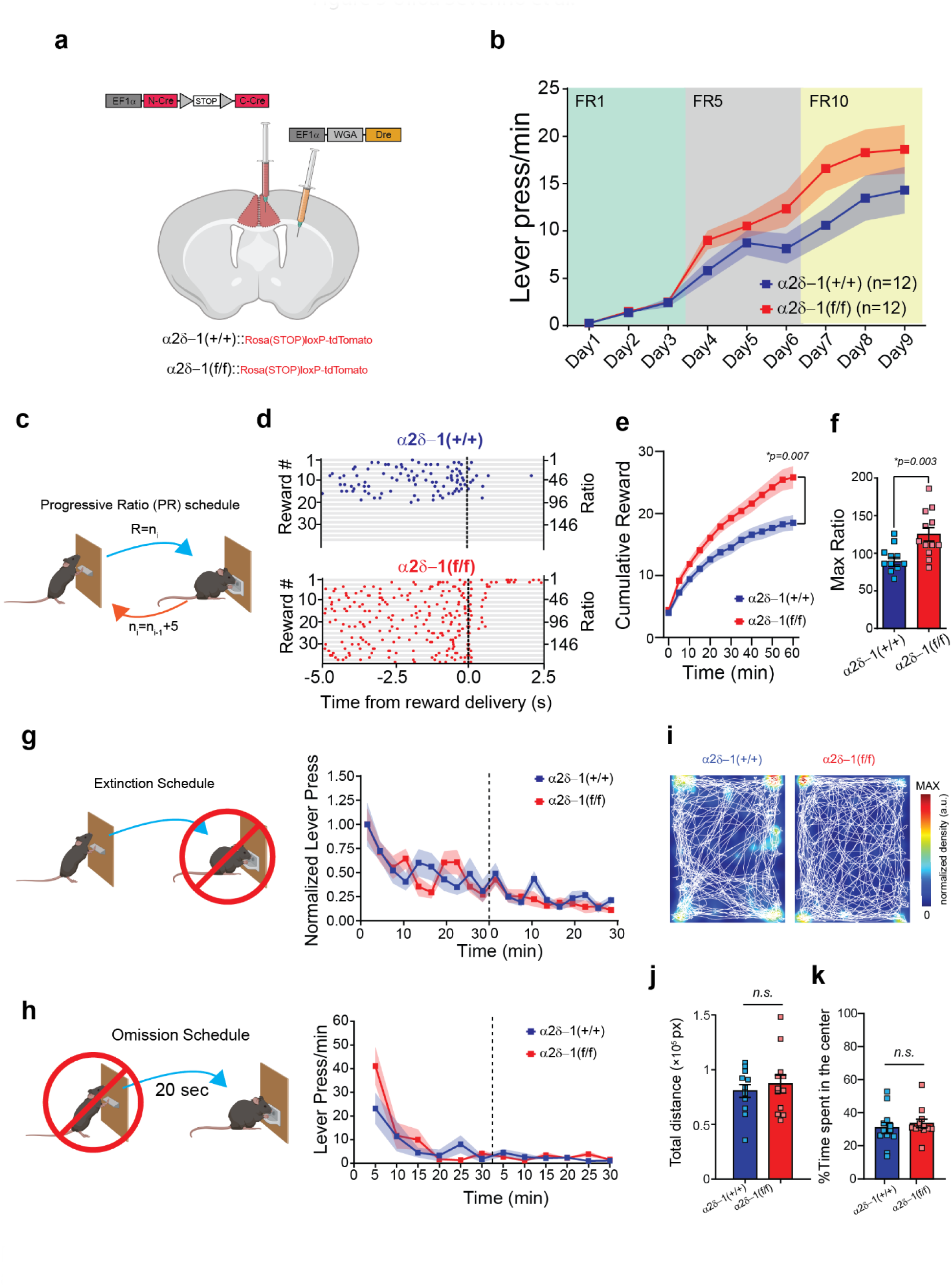
Conditional deletion of α2δ-1 from ACC_->DMS_ neurons increases effort exertion without affecting the learning of instrumental actions. a, Schematic representation of the injection strategy to label ACC_->DMS_ neurons and knock out α2δ-1. b, Lever press rate for the 9 days on the FR schedule for α2δ-1 (+/+) (n = 12; 6 male and 6 female) and α2δ- 1(f/f) (n = 12; 6 male and 6 female) mice. RM *Two-way ANOVA. Main effect of Days [F (1.795, 39.50) = 55.72, p < 0.0001], no effect of genotype [F (1, 22) = 2.851, p = 0.105] nor interaction [F (8, 176) = 1.943, p = 0.056].* c, Schematic representation of PR schedule. d, Representative peri-reward raster histograms of LP for α2δ-1 (+/+) and α2δ-1(f/f) mice. e, Cumulative reward count over the PR session (bin=5min) for α2δ-1 (+/+) (n = 12; 18.5 ± 1.2 rewards) and α2δ-1(f/f) animals (n = 12; 25.8 ± 1.8 rewards). *Two-way ANOVA for repeated measures. Main effects of Genotype [F (1, 22) = 8.832] and Time [F (12, 264) = 298.4, p < 0.0001] and interaction [F (12, 264) = 10.62, p < 0.0001].* f, Max Ratio for α2δ-1 (+/+) (n = 12; Max Ratio = 88.5 ± 6.1) and α2δ-1(f/f) (n = 12; Max Ratio = 125.2 ± 9.0) animals. *Unpaired t-test [t (22) = 3.366, p=0.003].* g, *left:* Schematic representation of the Extinction schedule in which mice do not receive any rewards upon lever press. *right:* Normalized lever press number in a 3 min bins for α2δ-1 (+/+) (n =12) and α2δ-1(f/f) (n = 12) animals. *Two-way ANOVA for repeated measures. Main effects of Time [F (5.433, 119.5) = 13.14, p < 0.0001], no effect of genotype [F (1, 22) = 0.1059, p = 0.748] nor interaction [F (19, 418) = 1.399, p = 0.122].* h, *left:* Schematic representation of the omission schedule for α2δ-1 (+/+) (n = 9) and for α2δ-1 (f/f) (n = 11). *right:* The lever press/min are reported for the 2 days of testing (dashed line). *RM Two-way ANOVA. Main effect of time [F (2.269, 40.85) = 15.81, p < 0.0001], no effect of genotype [F (1, 18) = 0.7326, p = 0.403] and significant interaction [F (11, 198) = 1.893, p = 0.042]. Multiple comparison showed no differences between genotypes at any time point.* i, Examples of open field patterns of locomotion by α2δ-1 (+/+) and α2δ-1 (f/f) mice. j, Bar graph of the total distance traveled in pixels for α2δ-1 (+/+) (n = 12; 8.0*10^4^ ± 5.8*10^3^ pixels) and α2δ-1(f/f) (n = 12; 8.7*10^4^ ± 8.2*10^3^ pixels) mice during the open field test. *Unpaired t-test [t (22) = 0.62, p = 0.538]*. k, Percent time spent in the center of the arena by α2δ-1 (+/+) (n = 12; 31 ± 3.3%) and α2δ-1(f/f) (n = 12; 33 ± 2.6%) mice. *Unpaired t- test [t (22) = 0.64, p = 0.526].* For all graphs: *Multiple comparisons using Holm-Sidak method; alpha = 0.05 for adjusted p-value*. Data shown as mean ± s.e.m.

Next, we tested the same mice on the PR schedule (Fig. 5c) and observed stark differences between α2δ-1(f/f) and α2δ-1(+/+) in effort exertion. The representative LP raster plots from two PR sessions, one from an α2δ-1(+/+) and the other from an α2δ-1(f/f) mouse, illustrate some of these differences (Fig. 5d). Similar to the α2δ-1 KO mice, α2δ-1(f/f) received a higher number of rewards across the session time (60 min) and reached a significantly higher maximum ratio than the α2δ-1(+/+) mice (Fig. 5e-f). Quantitative bout analyses revealed that the change in the performance of the LP sequence (Supplementary Fig. 5e-h) underlies the differences between genotypes. In α2δ-1(f/f) mice, bouts are longer and have a higher number of presses per bout, hence a higher frequency (lower bout IPI), than the α2δ-1(+/+) (#of presses/bout for α2δ-1(+/+) = 12 ± 1.7; and for α2δ-1(f/f) = 25 ± 3.4). These results show that ablating α2δ-1 specifically in ACC_->DMS_ neurons causes a profound alteration of the lever press sequence, resulting in increased effort exertion. On the other hand, deletion of α2δ-1 from ACC_->DMS_ neurons did not affect the ability of α2δ-1(f/f) mice to extinguish LP behavior, when the reward is no longer delivered (Fig. 5g) or to learn the opposite contingency (Fig. 5h). α2δ-1(f/f) mice also did not show general hyperactivity, as indicated by the open field test (Fig. 5i): there were no differences in total distance traveled (Fig. 5j) and time spent in the center of the arena (Fig. 5k) compared to α2δ-1(+/+). Altogether, these data show that loss of α2δ-1 only in the ACC_->DMS_ neurons is sufficient to cause a profound increase in effort exertion, when the task progressively becomes more demanding. Furthermore, these results suggest that α2δ-1 is required in the ACC_->DMS_ neurons for new excitatory synapse formation during LP learning to monitor action sequence and regulate effort exertion.

### Conditional deletion of **α**2**δ**-1 reduces the number and activity of excitatory synapses in the adult ACC_->DMS_ neurons

Why does the loss of α2δ-1 in the ACC_->DMS_ neurons cause an increase in effort exertion? Previously, dorsal-root ganglia neurons lacking α2δ-1 were shown to have reduced action potential (AP) firing frequency^87^ so, we wondered if changes in the AP firing frequency of ACC_->DMS_ neurons underlie this behavioral phenotype. To test this possibility, we performed current step stimulation of the Cre^+^ (tdTomato^+^) ACC_->DMS_ neurons of α2δ-1(+/+) and α2δ-1(f/f) mice (Fig. 6a, b) during whole-cell patch-clamp recordings. We observed no differences between the mean AP firing frequencies of the tdTomato^+^ neurons between the genotypes in any of the step current stimulations used to elicit neuronal APs (Fig. 6b, c). Moreover, we did not observe a difference in the mean resting membrane potentials between α2δ-1(+/+) and α2δ-1(f/f) neurons (Fig. 6d), showing that loss of α2δ-1 does not alter the excitability of the ACC_->DMS_ neurons.

**Figure 6.**
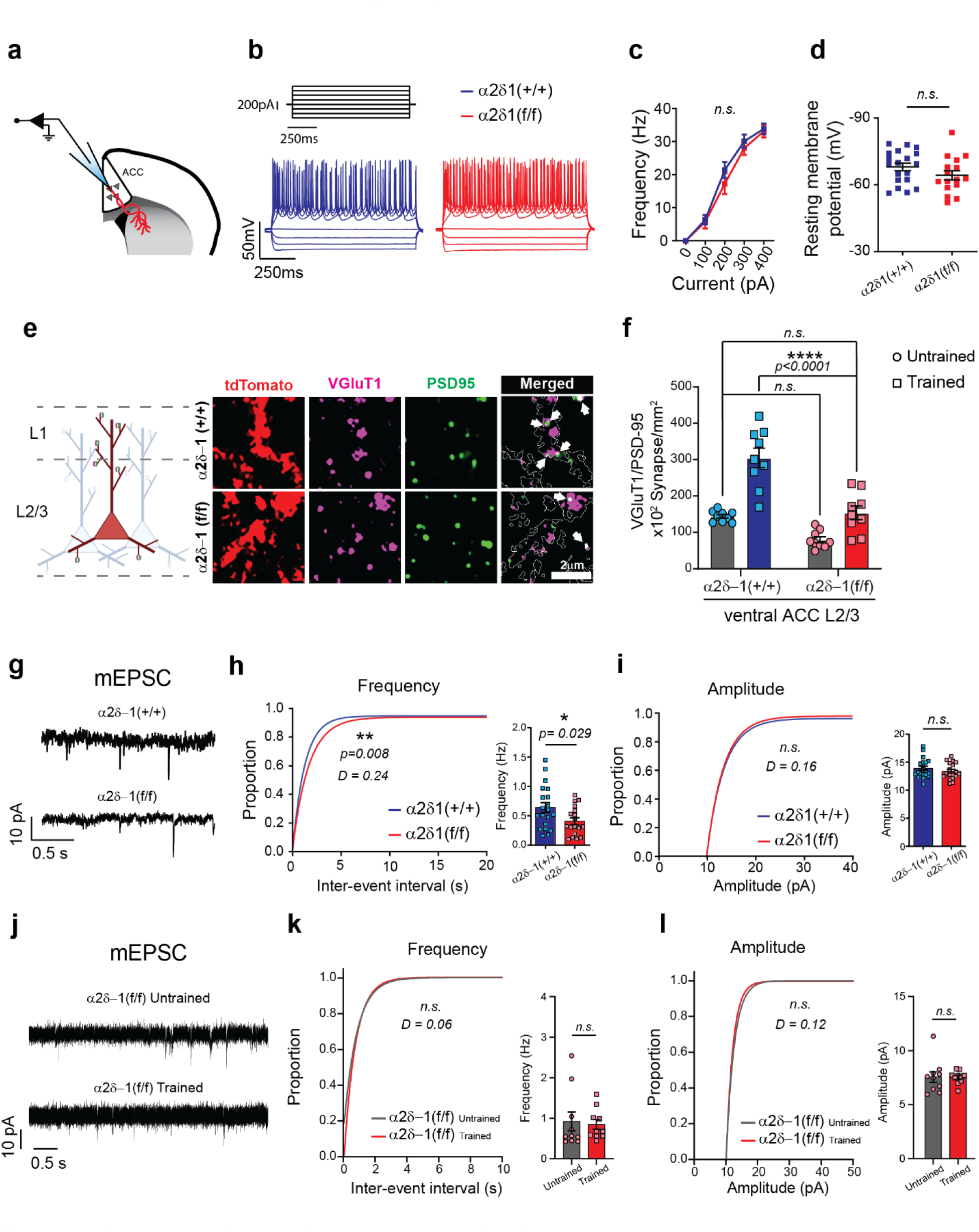
Circuit-specific conditional deletion of α2δ-1 reduces the excitatory synapses number onto the ACC_->DMS_ neurons. **a**, Schematic representation of the electrophysiological recordings from tdTomato^+^ neurons in both α2δ- 1(+/+) and α2δ-1(f/f) mice. **b**, Example traces of the intrinsic excitability test of α2δ-1(+/+) and α2δ-1(f/f) tdTomato^+^ neurons. **c**, Action potential (AP) frequency as function of the injected current for both α2δ- 1(+/+) (n = 4 mice, 19 cells) and α2δ-1(f/f) (n = 4 mice, 16 cells) animals. RM *Two-way ANOVA. Main effect of Current [F (2.3, 75.91) = 221.6, p < 0.0001], no effect of genotype [F (1, 33) = 0.5398, p = 0.467] and no interaction [F (4, 132) = 0.7386, p = 0.567].* **d**, Resting membrane potentials of tdTomato^+^ neurons in both α2δ-1(+/+) (n = 4 mice, 19 cells) and α2δ-1(f/f) (n = 4 mice, 16 cells) animals. *Unpaired t-test, [t (33) = 1.416, p = 0.166].* **e**, *left:* Schematic representation of the methodology used to quantify the VGluT1-PSD95 co-localized puncta. r*ight:* Representative images of the tdTomato positive neurons labeled for both VGluT1 and PSD95. The merged channel only shows the puncta within the tdTomato mask (*white arrows*). **f**, Comparison between vACC L2/3 untrained and trained α2δ-1(+/+) and α2δ-1(f/f) mice showing the VGluT1/PSD95-positive synaptic density. *Two-way ANOVA. Main effect of genotype [*F (1, 32) = 39.73, p < 0.0001*], Training [*F (1, 32) = 46.97, p < 0.0001*] and interaction [*F (1, 32) = 6.473, p = 0.016*]. Multiple comparison showed in figure for relevant comparisons.* n = 3 mice per condition and genotype (n = 3 images per mouse). **g**, Example traces from mEPSC recordings from trained α2δ-1(+/+) and α2δ-1(f/f) mice. **h**, *left:* cumulative distribution of the Inter-event interval of mEPSC in α2δ-1(+/+) (n = 4 mice; 20 cells) and α2δ-1(f/f) (n = 3mice; 18 cells) mice. *Kolmogorov-Smirnov test. right:* Average frequency of mEPSC in α2δ-1(+/+) (n = 4 mice, 20 cells; 0.64 ± 0.08 Hz) and α2δ-1(f/f) (n = 3mice, 18 cells; 0.41 ± 0.06 Hz) mice. *Unpaired t-test [t (36) = 2.27].* **i**, *left:* cumulative distribution of amplitude of mEPSC in pA for both α2δ-1(+/+) (n = 4 mice; 20 cells) and α2δ-1(f/f) (n = 3mice; 18 cells) mice. *Kolmogorov-Smirnov test [p = 0.176]. right:* Average amplitude of mEPSC in α2δ-1(+/+) (n = 4 mice, 20 cells; 14 ± 0.38 pA) and α2δ-1(f/f) (n = 3 mice, 18 cells; 13 ± 0.34 pA) mice. *Unpaired t-test [t (36) = 1.00, p = 0.322].* **j**, Representative traces from mEPSC recording from untrained and trained α2δ-1(f/f) mice. **k**, *left:* cumulative distribution of the Inter-event interval of mEPSC in untrained (n = 10 cells from 4 mice) and trained (n = 10 cells from 4 mice) α2δ-1(f/f) mice. *Kolmogorov-Smirnov test (p = 0.723).* Average frequency of mEPSC in Untrained (0.92 ± 0.23) and Trained (0.84 ± 0.12) α2δ-1(f/f) mice. *Unpaired Two- tailed t-test [t (18) = 0.296].* **l**, *left:* cumulative distribution of amplitude of mEPSC in untrained (n = 10 cells from 4 mice) and trained (n = 10 cells from 4 mice) α2δ-1(f/f) mice. *Kolmogorov-Smirnov test (p = 0.705). right:* Average amplitude of mEPSC in Untrained (7.6 ± 0.49) and Trained (7.5 ± 0.19) α2δ-1(f/f) mice. *Unpaired t-test [t (18) = 0.064, p = 0.949].* For all graphs: Data shown as mean ± s.e.m. alpha = 0.05.

In the visual cortices of constitutive α2δ-1 KO mice, L2/3 neurons displayed a severe reduction in the frequency of mEPSC compared to littermate WTs^59^, so we next tested whether loss of α2δ-1 in ACC_->DMS_ neurons would lead to a reduction in the density of synaptic inputs made onto these cells. To do so, first we quantified the number of VGluT1/PSD95-positive synapses made onto the tdTomato^+^ neuronal processes (Fig. 6e). These analyses were made both in L1 and L2/3 because the L2/3 ACC_->DMS_ pyramidal neurons extend their apical dendrites to L1 and basal dendrites to L2/3 (Fig. 6e). Training induced a significant increase in the density of VGluT1/PSD95-positive synapses made onto the α2δ-1(+/+) ACC_->DMS_ neuron processes in L1 (Supplementary Fig. 6a). Deletion of α2δ-1 from the ACC_->DMS_ neurons abolished the training-induced increase in synapse density in L1 (Supplementary Fig. 6b). However, in L2/3, training still induced a significant increase in the density of synapses in both genotypes (Supplementary Fig. 6a, b). However, the synapse density of trained α2δ-1(f/f) ACC_->DMS_ neurons was significantly lower than the trained α2δ-1(+/+) and not different from the untrained α2δ-1(+/+) (Fig. 6f). We recorded mEPSC from the ACC_->DMS_ neurons of trained α2δ-1(f/f) mice and trained α2δ-1(+/+) and we found a significant reduction in the frequency but not the amplitude of the mEPSCs (Fig. 6g-i).

As opposed to the training-induced structural and functional increases in excitatory synapses in WT mice (Fig 2d, e and 2h-j), in the α2δ-1(f/f) ACC_->DMS_ neurons we found that the observed structural increase is not sufficient to drive a functional change. Recordings of mEPSCs in the α2δ-1(f/f) ACC_->DMS_ neurons from L2/3 neurons of trained and untrained mice revealed no differences in the frequency or amplitude of the mEPSCs after training (Fig. 6j-l).

Taken together, these electrophysiological and neuroanatomical analyses show that circuit-specific conditional deletion of α2δ-1 in the adult ACC_->DMS_ neurons reduces the number and function of excitatory synaptic inputs made onto these neurons without changing intrinsic excitability. These results suggest that training-induced excitatory synaptogenesis is required to properly excite the ACC_->DMS_ neurons and the activity of ACC_->DMS_ neurons control effort exertion by diminishing lever press behavior when the task becomes too demanding.

### Optogenetic excitation of ACC_->DMS_ neurons is sufficient to reduce effort exertion in WT and **α**2**δ**-1 circuit-specific knockout mice

To determine if the excitation of ACC_->DMS_ neurons is sufficient to reduce effort exertion during the LP behavior, we expressed a Cre-dependent light-gated cation-selective membrane channel, Channelrhodopsin-2 (flex-ChR2)^88^, in these neurons. To do so, α2δ-1(+/+) and α2δ- 1(f/f) mice received bi-lateral injections of the WGA-Dre virus in the DMS and a 1:1 cocktail of N-Cre-*rox-STOP-rox*-C-Cre and flex-ChR2 viruses in the ACC (Fig. 7a). Optic fibers were also implanted bilaterally in the ACC to enable activation of ACC_->DMS_ neurons (Fig. 7a). To compare the LP behavior of the same mouse with or without optogenetic stimulation of ACC_->DMS_ neurons, we tested the mice with a more demanding fixed ratio schedule FR20. For these optogenetic experiments we cannot use the progressive ratio schedule, because the LP/reward ratio changes over time. Thus, during PR schedule we cannot compare the lever press rate of the same mouse with or without stimulation within the same session.

**Figure 7.**
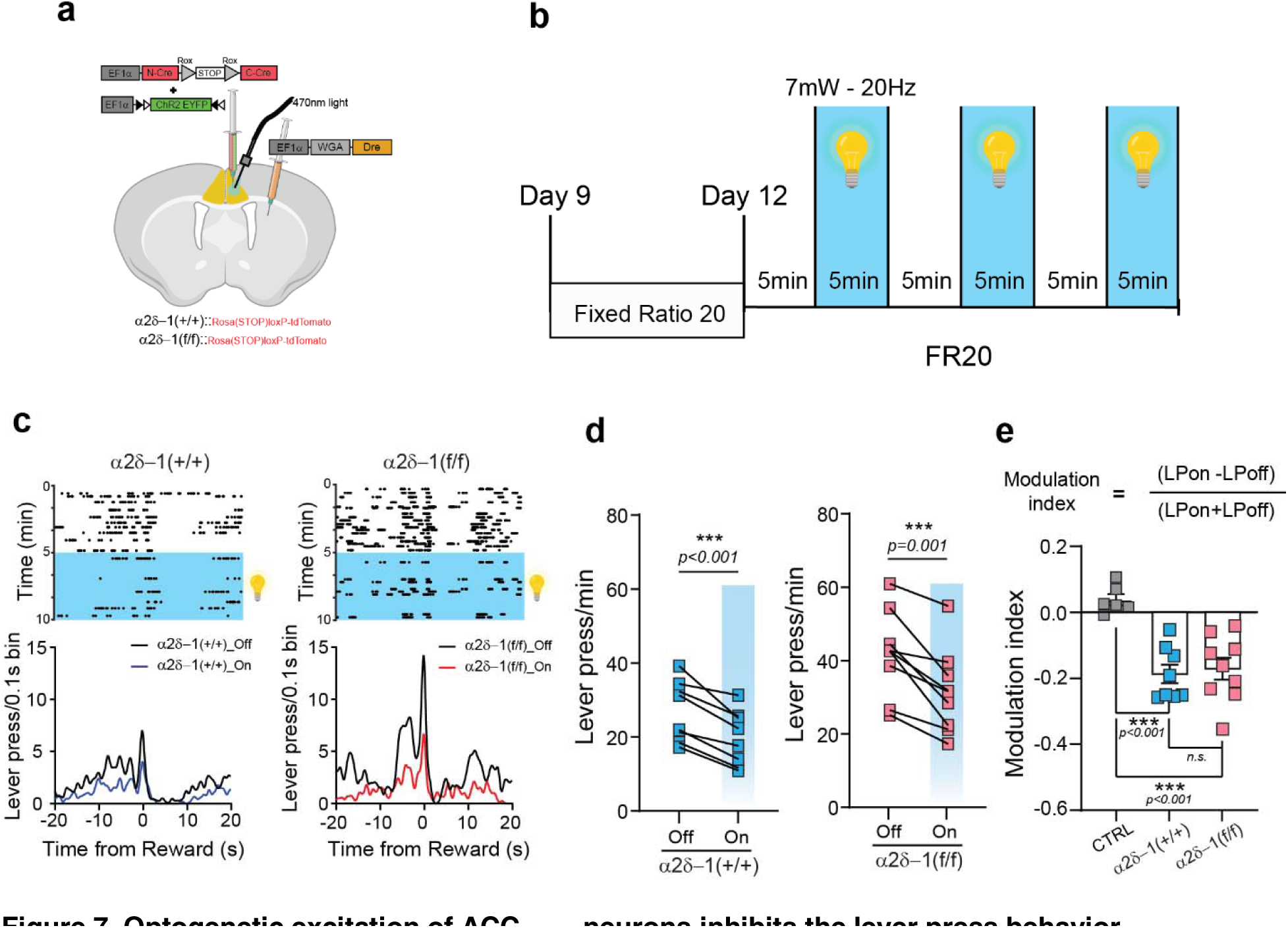
Optogenetic excitation of ACC_->DMS_ neurons inhibits the lever press behavior. **a**, Schematic representation of the viral injections and fiber implants for the optogenetic rescue experiments in the ACC_->DMS_ projecting neurons of α2δ-1(+/+) and α2δ-1(f/f) mice. **b**, Schematic representation of the FR20 schedule used for the optogenetic experiments. **c**, Representative peri-reward raster histograms of an α2δ-1(+/+) and an α2δ-1(f/f) mouse during the light-Off and light-On periods. **d**, Lever press/min for the α2δ-1(+/+) mice (n = 8) and α2δ-1(f/f) mice (n = 11). *Paired Two-tailed t-test* for α2δ-1(+/+) light-Off (27 ± 2.9) and light-On (20 ± 2.6) *[t (7) = 6.9]* and for α2δ-1(f/f) light-Off (42 ± 3.8) and light-On (32 ± 3.8) *[t (10) = 6.7].* **e**, Modulation index for CTRL, α2δ-1 (+/+), and α2δ-1 (f/f) mice. *One-way ANOVA [F (2, 20) = 16.27, p < 0.001], Tukey’s multiple comparison revealed a significant difference between CTRL and* α*2δ-1 (+/+) [q (20) = 7.3] and between CTRL and* α*2δ-1 (f/f) [q (20) = 7.0] and no differences between* α*2δ-1 (+/+) and* α*2δ-1 (f/f) [q (20) = 0.56, p = 0.916].* For all graphs: Data showed as mean ± s.e.m. alpha = 0.05.

Mice were trained for 9 days as described previously. Subsequently, they were trained for an additional two days on an FR20 schedule (Fig. 7b). On day 12, we tested these mice over a 30 min session, during which we alternated 5 minutes of optogenetic stimulation (light-On) with 5 minutes of no stimulation (light-Off) (Fig. 7b). In a control group of mice (CTRL), which had only fiber implants into the ACC but no virus, we controlled for possible effects of the light exposure and surgery on behavioral performance (Supplementary Fig. 7a)^89^. We observed that these CTRL mice behave similarly during periods of light-On and light-Off, showing that light exposure in the absence of ChR2 expression had no or little effect on their behavior (Supplementary Fig. 7b, c). For each mouse, we confirmed the anatomical locations of the optic fiber placement and the efficiency of viral targeting and co-expression of Cre (tdTomato^+^) and ChR2 (EYFP^+^) in the ACC_->DMS_ neurons (Supplementary Fig. 7d, e). The co-localization of the tdTomato (Cre) and EYFP (ChR2) signals was observed in more than half of the tdTomato^+^ cells (Supplementary Fig. 7f, g), with L2/3 having the highest percentage of positive cells among all the layers (Supplementary Fig. 7h).

As shown by the representative raster plots and histograms (Fig. 7h), in the absence of optogenetic stimulation, the numbers of LPs are increased in α2δ-1(f/f) compared to α2δ-1(+/+) mice (Fig. 7c, d). However, during the light-On intervals, both α2δ-1(+/+) and α2δ-1(f/f) mice reduced their LP rate (Fig. 7c, d). To quantify this change, we calculated the modulation index denoting the difference in the number of LPs per 15 minutes light-On compared to the 15 minutes of the light-Off periods divided by the total number of LPs for the entire 30-minute session (Fig. 7e). The mice expressing ChR2 in the ACC_->DMS_ neurons displayed a 20% reduction in the number of LPs when stimulated (Fig. 7e). Importantly, planned t-test analysis showed that under light-On conditions, the number of LP of the α2δ-1(f/f) mice were comparable with those of the α2δ-1(+/+) mice during light-Off periods (Fig. 7d, *planned unpaired two-tails t-test [t (15) = 0.98], p=0.344*). These results indicate that when α2δ-1 is deleted in ACC_->DMS_ neurons, LP behavior can be rescued by optogenetic activation of these neurons. Taken together, we found that optogenetic excitation of ACC_->DMS_ neurons is sufficient to reduce the effort exerted during a demanding task. Furthermore, when considered together with our anatomical and functional analysis (Fig. 6) these data strongly indicate that α2δ-1-mediated synaptogenesis is required for the ACC_->DMS_ neuronal activation necessary to regulate effort exertion and suggest that reduced activity of the ACC_->DMS_ neurons can result in an increase in the amount of effort exerted during task performance.

### Optogenetic inhibition or excitation of ACC_->DMS_ neurons inversely modulate the effort exerted during a demanding task

To test if inhibition of ACC_->DMS_ neurons could phenocopy the α2δ-1(f/f) mice and cause an increase in effort exertion, we expressed an inhibitory step-function opsin called BLINK2 in ACC_->DMS_ neurons (Fig. 8a). BLINK2 activation by light provides sustained neuronal inhibition after a short period of optogenetic stimulation ^90^. For these experiments, mice were tested on a PR schedule with and without optogenetic stimulation. We found that 1 minute of light delivery at the beginning of a PR schedule causes an increase in the number of lever presses (Fig. 8b). After 10 minutes, the inhibitory effects of the optogenetic manipulation subside, and the LP rate returns to the levels observed without stimulation (Fig. 8b). We statistically analyzed the effects of BLINK2-mediated inhibition during the first 10 minutes after stimulation onset. These data show that inhibition of the ACC_->DMS_ neurons is sufficient to increase LP rate, the number of rewards obtained, and the maximum ratio reached when compared to performance of the same mice on a day in which they did not receive optogenetic stimulation (Fig. 8c-e). Analysis of the lever press bouts revealed that inhibition of ACC_->DMS_ neurons also increases the frequency and number of lever presses per bout as observed in α2δ-1(f/f) mice (Fig. 8f-i).

**Figure 8.**
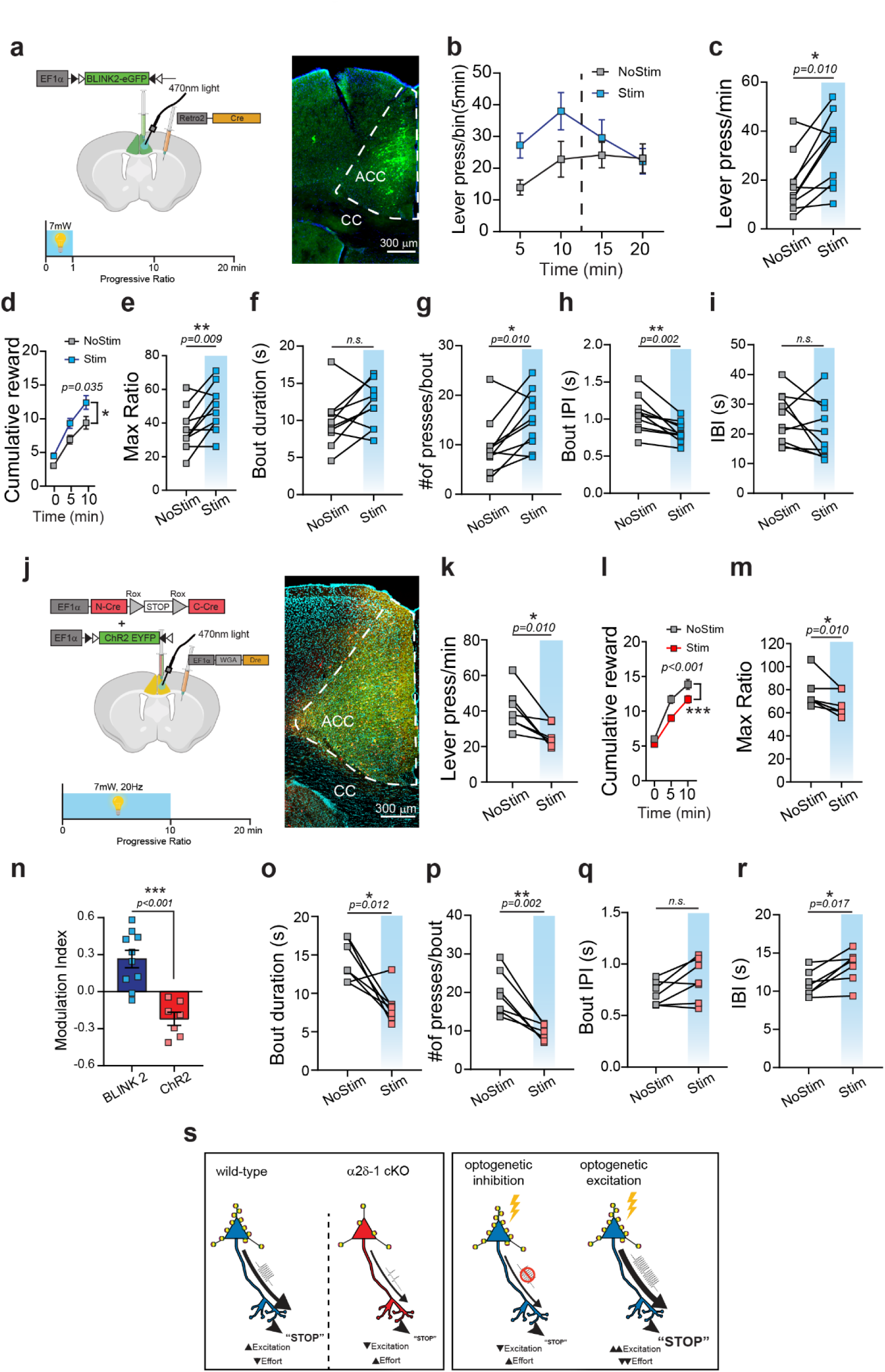
Inhibition and excitation of ACC_->DMS_ neurons have opposite effects on the performance of lever press bout and effort exertion. a, *left:* Schematic representation of viral injections and behavioral scheduled used for inhibitory experiments and *right:* example image of BLINK2 expression within ACC_->DMS_ neurons. b, Lever press/min of the entire PR test for BLINK2 mice showing the effect of BLINK2 inhibition for the first 10 minutes upon stimulation. c, Quantification of behavioral parameters during the first 10min of PR for not stimulated and stimulated conditions (n = 10 mice). Lever press/min during NoStim (18 ± 3.8) and Stim (30 ± 4.5) days of PR. *Paired Two-tailed t-test [t (9) = 3.2].* d, Cumulative reward count during NoStim and Stim days of PR. *RM Two-way ANOVA, main effect of stimulation [F (1, 18) = 5.219] and time [F (2, 36) = 161.1, p < 0.0001], no interaction [F (2, 36) = 0.6485, p = 0.528]*. e, Max Ratio between NoStim (37 ± 4.0) and Stim (49 ± 4.4) days. *Paired Two-tailed t-test [t (9) = 3.3]*. f, Bout duration in NoStim (9.8 ±1.1) and Stim (12 ± 0.96) conditions were not significantly different. *Paired Two-tailed t-test [t (9) = 2.0, p = 0.074].* g, Number of presses per bout during NoStim (9.7 ± 1.8) and Stim (15 ± 1.8) days showed a significant difference *Paired Two-tailed t-test [t (9) = 3.2].* h, Bout IPI during NoStim (1.1 ± 0.07) and Stim (0.8 ± 0.04) showed a significant decrease. *Paired Two-tailed t-test [t (9) = 4.2].* i, IBI in NoStim (25 ± 2.6) compared to Stim (21 ± 3.0) showed no difference. *Paired Two-tailed t-test [t (9) = 1.4, p = 0.186].* j, *left:* Injection strategy and protocol used for optogenetic experiments using ChR2 during PR schedule. *right:* example image of ChR2 expression in ACC_->DMS_ neurons. k, Quantification of behavioral parameters during the first 10min of PR for not stimulated and stimulated conditions (n = 7 mice). Lever press/min during NoStim (41 ± 4.4) and Stim (26 ± 2.4) days of PR. *Paired Two-tailed t-test [t (6) = 3.7].* l, Cumulative reward count during NoStim and Stim days of PR. *RM Two-way ANOVA, main effect of stimulation [F (1, 12) = 10.21] and time [F (1.458, 17.49) = 267.4, p < 0.0001], significant interaction [F (2, 24) = 5.379, p = 0.012]*. m, Max Ratio between NoStim (77 ± 5.2) and Stim (67 ± 3.8) days. *Paired Two-tailed t-test [t (6) = 3.2]*. n, Modulation index of BLINK2 (0.26 ± 0.07) and ChR2 (-0.22 ± 0.05) mice during the PR schedule. *Unpaired Two-tailed t-test [t (15) = 5.1].* o, Bout duration in NoStim (14 ±1.0) and Stim (8.3 ± 0.86) conditions were significantly different. *Paired Two-tailed t-test. [t (6) = 3.6].* p, Number of presses per bout during NoStim (20 ± 2.2) and Stim (9.7 ± 0.79) days showed a significant difference. *Paired Two-tailed t-test [t (6) = 5.0]*. q, Bout IPI during NoStim (0.71 ± 0.04) and Stim (0.85 ± 0.08) showed no significant difference*. Paired Two-tailed t-test [t (6) = 2.4, p = 0.051].* r, IBI in NoStim (11 ± 0.63) compared to Stim (13 ± 0.80) showed a significant difference. *Paired Two-tailed t-test [t (6) = 3.3].* s, *left:* Proposed intrinsic mechanism of regulation of effort exertion through training-induced excitatory synaptogenesis. *right:* representation of how optogenetic manipulation of ACC_->DMS_ neurons would affect the effort exertion. For all graphs: *Multiple comparisons using Holm-Sidak method; alpha = 0.05 for adjusted p-value*. Data shown as mean ± s.e.m.

Next we performed optogenetic excitation of ACC_->DMS_ neurons under similar conditions in mice co-expressing Cre (tdTomato^+^) and flex-ChR2 (EYFP^+^) in the ACC_->DMS_ neurons (Fig. 8j). In contrast to the inhibition of ACC_->DMS_ neurons, light stimulation during the first 10 minutes of a PR schedule caused a significant reduction in the number of LP, cumulative rewards and maximum ratio (Fig. 8k-m). Indeed, calculation of the modulation index for inhibition and excitation of ACC_->DMS_ neurons shows opposite effects of optogenetic manipulation on mouse behavior (Fig. 8n). Analysis of the lever press bouts revealed that the reduced effort exertion was due to the performance of shorter bouts with lower number of presses (Fig. 8o-r).

These results reveal that the reduced or increased activity of ACC_->DMS_ neurons regulate the amount of effort exerted during a demanding task by modulating the lever press bout properties. These data also suggest a role for the ACC_->DMS_ neurons in monitoring and controlling the performance of action sequence during goal-directed actions.

## Discussion

Synaptogenesis is critical for the construction of brain circuits during development ^5, 7, 62, 91, 92^. However, little is known about the role of synaptogenesis in the adult brains. Here we show that new excitatory synapses are formed onto the ACC_->DMS_ neurons during instrumental operant conditioning training. These new synapses are not necessary for learning the action-outcome relationship but are required for adjusting the effort in an adaptive manner. The Thrombospondin-Gabapentin receptor α2δ-1 mediates training-induced excitatory synapse formation onto the ACC_->DMS_ neurons. When α2δ-1 is lost, training-induced excitatory synaptogenesis is diminished, generating a maladaptive behavioral phenotype in which mice spend significantly more effort for achieving the reward. Concordantly, circuit-specific excitation of ACC_->DMS_ neurons is sufficient to reduce the effort exerted during instrumental actions; whereas their inhibition increases the effort exerted. Taken together, our findings offer a circuit-specific cellular and molecular mechanism for cognitive control of effort (Fig 8s).

In our behavioral paradigm, the number of lever presses represent the cost, whereas the food pellet reward is the benefit. When the task becomes too demanding (e.g., in PR or FR20 schedules), WT animals reduce the amount of effort because the cost of reward gets too high. Our findings show that loss of α2δ-1-signalling disrupts this adaptive step. We used the PR test to measure effort exertion, because it avoids the confound of having to select between different options, such as choosing between two actions with different probability of reward or two different locations with different rewards ^20, 38, 40, 68, 93, 94^. In fact, PR test is a standard test to measure effort exertion both in rodents and humans^37, 84–86^.

To interpret the behavioral phenotype in α2δ-1 global and circuit-specific KOs as “increased effort exertion”, we ruled out other possibilities. A possible alternative explanation for increased lever press behavior in α2δ-1 KOs could be the establishment of persistent or habitual behaviors. If that were the case, we would expect to see persistent lever pressing during the extinction test or no effect on lever pressing after outcome devaluation. But instead, when we performed extinction and devaluation tests, the KO mice were able to extinguish the behavior and adjust their effort after reward devaluation (Fig. 4, S4, and 5). Moreover, mice with α2δ-1 deletions did not show impaired reversal learning, further showing that learning the action-outcome relationship is intact in α2δ-1 KOs (Fig. 4 and 5). Finally, general hyperactivity was ruled out using an open field test to evaluate the overall mouse movement in a novel environment (Fig. 4 and 5). Moreover, optogenetic excitation or inhibition of only the ACC_->DMS_ neurons is sufficient to modulate lever press rates within the same mice, showing that this particular circuit is a regulator of effort exertion (Fig. 7 and 8).

Loss of α2δ-1 or optogenetic modulation of ACC_->DMS_ neurons cause increased effort exertion by affecting the action sequence (bout) properties. Particularly, as the effort exertion increase, the number of lever presses per bout increased and the inter-press interval decreased. These changes occurred without an increase in bout duration or inter-bout intervals (Fig. S5, and 8). These results show that the ACC_->DMS_ neurons control effort exertion by modifying the rate of lever press within a bout.

In mice and non-human primates, ACC activity is proposed to be necessary during behavioral tasks in which past or current action-outcome contingencies guide future behavioral decisions ^95–97^. For example, in a foraging simulation, monkeys with ACC lesions were more likely to repeat a learned action even if the probability of receiving the reward decreased^96^, suggesting that ACC function is involved in evaluating the changes in the relationship between the effort and reward. The time scale of synaptogenesis (days) is unlikely to underlie the online adjustment of performance by ACC. Instead, our findings suggest that new synapses are formed onto the ACC_->DMS_ neurons during training as an anticipatory mechanism for future scenarios in which the effort needed for the learned behavior increases.

New findings suggest that training-induced modification of synaptic connectivity could be a common mechanism used by other PFC regions to guide an adaptation of behavioral response. For example, learning induces the formation of new spines in the Orbitofrontal cortex (OFC) neurons projecting to the DMS. These newly formed spines are necessary to retrieve previous memories of learned actions and select actions with higher reward^20^. It is possible that OFC and ACC connections to the DMS have synergistic functions to control adaptation of goal-directed learned behaviors.

Previously, we found that the Thrombospondin-Gabapentin receptor, α2δ-1, is required for the formation of intracortical excitatory synapses in the developing visual cortex^59^. Loss of α2δ-1 causes more than 50% reduction in excitatory synapse numbers, function, and dendritic spines. Here, we found that α2δ-1 is required for proper intracortical connectivity in the adult ACC as well. The lack of α2δ-1 is sufficient to abolish the training-induced increase in excitatory synapse numbers and activity in this brain region (Fig. 2,3, and 6).

New synapse formation in the motor cortex was proposed as an important step in motor learning and memory^98, 99^. Thus, initially, we expected the α2δ-1 mutants to have deficits in the learning phase of the instrumental training. Another study, using a different α2δ-1 knockout line and behavioral paradigms, reported that lack of α2δ-1 impairs motor and spatial learning^100^. Surprisingly, we found no learning impairments in α2δ-1 global or circuit-specific KO mice in instrumental training. Instead, our findings show that α2δ-1-signaling in adult controls effort exertion by regulating the formation of excitatory synapses onto the ACC_->DMS_ neurons.

α2δ-1 is a neuronal receptor for Thrombospondins (TSPs), which are astrocyte-secreted synaptogenic proteins. TSP-α2δ-1 signaling stimulates the formation of silent, structural synapses (e.g., NMDA receptors), whereby astrocytes can then activate them by recruiting AMPA receptors to the synapse^101^. Our findings imply that astrocyte-to-neuron signaling might be upstream of training-induced synaptogenesis. Future studies investigating the roles of astrocytes and the TSPs that they secrete in the control of goal-directed learned behaviors might be fruitful.

The importance of synaptogenesis in brain development is well-established; however, whether synaptogenic signaling performs critical functions in the adult brain to regulate complex, learned behaviors is unknown. Mutations in genes controlling synapse formation and maturation are strongly linked to many neuropsychiatric disorders, including ASD, Schizophrenia, OCD, and Alzheimer’s Disease ^42, 43, 102–104^. A common hallmark of these disorders is the presence of repetitive and ineffective behaviors^105, 106^. Our findings reveal a critical link between synaptogenic signaling in the adult brain and the control of the adaptability of learned behaviors, suggesting that dysfunctional adult synaptogenesis may underlie the etiology of behavioral inflexibility seen in such disorders.

## Acknowledgments

This work was supported by grants from the National Institutes of Health (NS096352 and AG059409 to CE and MH112883 and DA040701 to HHY). FPUS was supported by postdoctoral fellowships from the Regeneration Next Initiative. Illustrations were created with BioRender.com. We would like to thank: Donna Porter, for helping with mouse colony maintenance; William Wetsel, Stacy Bilbo, Dolores Irala, Dhanesh Sivadasan Bindu, Alessandro De Simone and Sehwon Koh for their critical feedback on the manuscript. CE is an HHMI Investigator.

## Author Contributions

Conceptualization, FPUS, HY, and CE; Methodology, FPUS, OL, KS, NK, AF, SJ, SW, RH, HY, and CE; Investigation, FPUS, OL, KS, SJ, SW, NK and RH; Formal analysis, FPUS, OL, KS, SW, NK, and CS; Resources, IHK; and SS; Writing – original draft, FPUS, HY, and CE; Writing – Review & Editing, FPUS, OL, KS, HY, and CE Funding Acquisition, FPUS, HY, and CE.

## METHODS

### Animals, housing, and genotyping

All mice were used in accordance with the Institutional Animal Care and Use Committee (IACUC) and the Duke Division of Laboratory Animal Resources (DLAR) oversight (IACUC Protocol Numbers A173-14-07, A147-17-06, and A263-16-12). All mice were housed under typical day/night conditions of 12-hours cycles. Wild type C57BL/6J (Stock #000664) and ROSA(STOP)loxP-td-Tomato (B6;129S6-Gt(ROSA)26Sor^tm^^14(CAG-tdTomato)^Hze/J; Stock #007914) line were obtained through Jackson. The constitutive α2δ-1 Het and KO mice, as well as the conditional α2δ-1(f/f) mouse lines, were generated from our laboratory ^59^. All male and female mice (3-5 months old) used in this study were handled for 5-10 min a day for a week to allow them to get used to the operator. After this time, animals were food-restricted for 3-5 days until they rich 85-90% of their normal body weight. The target weight was maintained stable by daily feeding them, with 1.5/2 g of home chow, after training.

### Instrumental training

Lever pressing training was performed in operant chambers (St Albans, VT, USA) set within light resistant and sound attenuating walls. Each chamber was equipped with a food magazine, where an infrared beam recorded the head entries into the magazine, that received pellets from a dispenser. The delivered reward was a Bio-Serv 14mg Dustless Precision Pellets (Bio-Serv, NJ, USA). Each chamber also had two retractable levers on either side of the magazine and a 3-W 24-V house light mounted to the opposite side of the levers. A computer with the Med-PC-IV program was able to control the equipment and record the behavior according to the desired schedule. Timestamps for each lever press and head entries were recorded with a 10ms resolution and then analyzed using custom-written programs (available upon request). Lever pressing training was performed using only the left lever.

#### Fixed Ratio schedule

To test the capability of learning new behaviors, we used a continuous reinforcement schedule, here named Fixed Ratio 1 (FR1). The first day of FR1 began with 3 food pellets left in the food magazine, allowing the mice to learn about the possibility to receive food and the location of food delivery, the left lever inserted and the house light illuminated. The initial schedule consisted of 3 days of FR1, during which the animals received a pellet for each lever press. The session ended when one of the two restrictions was reached, 120 min or 50 rewards, with the retraction of the lever and the light turned off. At the end of the 3 days of FR1, mice were moved to 3 different testing schedules. Three days of FR5 schedule, during which the animals received a pellet for every 5 lever presses, were followed by the other three days of FR10 increasing the number of lever presses up to 10 for one reward. Session ended when one of the two restrictions was reached, 60 min or 50 rewards, with the retraction of the lever and the light turned off. The lever press bout analysis was performed using Matlab. The average inter press interval probability distribution was used to define the bout start and bout end events. In all the cases in which multiple conditions were compared, the control condition was used to define bout start and bout end events for both groups to identify changes from the control condition. At the end of these 9 days, the mice were either sacrificed within 1 hour for histological analysis and RNA purification or shifted to the next step of the behavioral test.

#### Progressive Ratio, Extinction, Omission and Revaluation tests

A Progressive Ratio (PR) schedule was used to evaluate the motivation of the mice. The PR schedule consists of an increasing number of lever presses every time a reward is received. In our case, we decided to use 5 as increased progression. The session ended after 60 minutes with the retraction of the lever and the light turned off. Extinction schedule occurred across two consecutive days during which mice could press the lever but there was no food delivered as a reward. The session ended after 30 minutes with the retraction of the lever and the light turned off. Omission schedule occurred across two consecutive days during which the reward was delivered at a fix interval of 20 secs unless a lever press happened. At every pressed performed by the mouse, the timer for the reward delivery reset resulting in a delay in the reward delivery. Outcome devaluation was performed after training the mice up to FR10 schedule. On one day mice were exposed to 0.5 g of home cage food for 30min (valued state). After retraining for FR10, mice were then exposed to 0.5 g of reward pellets for 30min (devalued state). After each pre-feeding session, a 10 min extinction test was given. The test began with the illumination of the house light and insertion of the left lever, and ended with the retraction of the lever and the offset of the house light. The number of presses on each lever was recorded.

### Open field test

After the operant task under food deprivation was completed, mice were normally housed with food and water provided ad libitum for 2-3 days, until their normal body weight was reestablished. Mice were then tested in an open field chamber equipped with a blackfly camera (Flyr System, BFS-U3-04S2M-CS) to record the animal movement. The Bonsai software (https://bonsai-rx.org/) was used to automatically detect and record the x and y coordinates for the center of mass of the mouse. The session began, after the mice were acclimated for 30min to the new room. The mouse was positioned to the center of the arena at the start of the recording which then finished after 30min. The chamber was carefully cleaned and the bedding was changed between different groups of littermates to avoid distraction due to other animals’ odor stimuli.

### c-Fos staining and analysis

Both trained WT mice (n=6) and untrained WT mice (n=6) were anesthetized with 200 mg/kg tribromoethanol (avertin) and then terminated by perfusing with a solution made of TBS with heparin (0.1128g Heparin ammonium salt from porcine intestinal mucosa [Sigma; H6279]) and then with 4% Paraformaldehyde (PFA) within 1 h after the last day of FR10 schedule to observe changes in the activity-dependent expression of the immediate-early gene c-Fos. Mouse brains were then kept in 4% PFA o.n. at 4°C. The day after brains were rinsed 3 times with TBS, immersed in 30% Sucrose in TBS and stored at 4°C until they were not floating anymore. At this time brains were included in a mixture of 30% Sucrose in TBS, and Tissue Tek O.C.T. compound (frozen tissue matrix) at a 1:2 ratio and stored at -80°C. Using a cryostat, the brains were cut into 25-30 μm coronal sections and stored floating in 50% Glycerol in TBS in a 24 multiwell plate, 5 sections per well, to have a 100 μm spatial representation in the rostrocaudal direction. 4 sections for each mouse were selected at these approximate (± 0.2) coordinates relative to Bregma: +2.2, +1.7, +0.4 and -0.3. The sections were rinsed 3 times (1x2min, 1x20min, and 1x30min) in TBS, then they were moved for 1h in a blocking solution containing 5% Normal Donkey Serum (NDS) in TBS+0.3% v/v of Triton. Primary antibody, Rabbit anti-cFos (Calbiochem, PC05), was diluted 1:50 in the blocking solution + 0.1% v/v of Sodium azide. The sections were incubated in this solution for 48-72h at 4°C with gentle shaking. After this time, they were transferred to a secondary antibody solution after 3 washing steps in TBST (Triton 0.3%) (1x10min, 1x30min, 1x40min). Donkey anti-rabbit conjugated to Alexa fluor-594 was used as secondary Ab in blocking solution to incubate the sections 2h at RT. DAPI (1:50000) was added 15 min before the end of this step and then sections were rinsed again 3 times as before mounting them on a Superfrost Plus slides using a Refractive Index solution (RI solution: 20mM Tris (pH 8.0), 0.5% N-propylgallate, 90% Glycerol).

Tile scan images were then acquired using an Olympus Fluoview confocal microscope using 20X lens in resonant scanner mode, allowing the fast acquisition of entire coronal sections with an optical section step of 1 um and the necessary number of images/stacks to acquire the entire depth of the section. Images were then processed using a combination of software. First, the images were restored using Content-aware image restoration (CARE) ^107^, this neural network allowed us to use images (minimum of 30 images used to train the code) acquired in galvo scanner mode for the restoration of the tile scan acquired in resonant scanner mode. The only requirement is that for both the scanner modes the same parameters have to be used for the acquisition (i.e. type of objective and zoom). Then flat field correction and stitching were performed with a custom code (https://github.com/ErogluLab/CellCounts) and finally, segmentation for DAPI^+^ and c-Fos^+^ (from now called c-Fos^+^) cells was performed using the UNet software (https://arxiv.org/abs/1505.04597, and adapted by Chaichontat (Richard) Sriworarat). In this case, images that were manually segmented by the user to indicate the DAPI^+^ and c-Fos^+^ cells, as well as ROIs that were not positive for either marker, were used to train the UNet neural network. The training process was reiterated until the segmentation was able to recognize the positive cells correctly on a small set of data and then applied to analyze the entire batch of images. The segmentation step produced 16-bit images with the mask of the segmented c-Fos^+^ cells in which the intensity is indicative of the degree of confidence of the segmentation. The mask was then used to select, based on the intensity, and count the c-Fos^+^ cells using the WholeBrain software ^67^. All the brain regions represented by less than 3 mice were not considered in the analysis.

### RNA sequencing preparation and analysis

RNA-sequencing libraries were made from ≥500ng of purified mouse ACC (n = 6 mice per condition; n = 3 mice per sex; 1 untrained male mouse was removed from the final analysis for having a low number of reads) RNA using the Kapa Stranded mRNA-seq kit. For each replicate 40-72 million, 2x51 reads were obtained from a NovaSeq 6000. Raw reads were adapter trimmed using Trimmomatic ^108^ (v0.38), aligned to the reference mouse genome (mm10) using Bowtie2 ^109^ (v2.3.5.1), and counted using Subread ^110^ (featureCounts ^111^, v1.6.3). Differential gene expression was conducted using edgeR ^112^ (v3.30.3). RNA-sequencing data have been deposited in the Gene Expression Omnibus (GEO) repository with accession number: <u>GSE169392</u>.

### Immunohistochemistry

Brain sections were washed three times then permeabilized in TBS with 0.2% Triton-X 100 (TBST; Roche, Switzerland) at room temperature. Sections were blocked in 5% Normal Goat Serum (NGS) in TBST for 1 h at room temperature. Primary antibodies (guinea pig anti-VGlut1 1:2000 [AB5905, Millipore, MA], rabbit anti-PSD95 1:300 [51–6900, Invitrogen, CA], guinea pig anti-VGAT 1:1000 [Synaptic Systems 131 004], rabbit anti-Gephyrin 1:500 [Synaptic Systems 147 002], were diluted in 5% NGS containing TBST. Sections were incubated overnight at 4°C with primary antibodies. Secondary Alexa-fluorophore (488, 594, and 647) conjugated antibodies (Invitrogen) were added (1:200 in TBST with 5% NGS) for 2 h at room temperature. Slides were mounted in Vectashield with DAPI (Vector Laboratories, CA) and images were acquired on an Olympus Fluoview confocal microscopy using a 60X oil lens at 1.64X Zoom.

### Quantification of Synapses

3-4 animals per genotype of WT, α2δ-1 KO, α2δ-1(f/f), and α2δ-1(+/+) were used for synapse analysis. Three independent brain sections per group were used for immunohistochemistry. 5 µm thick confocal z-stacks (optical section depth 0.33 µm, 15 sections/z-stack) of the ACC (layer 1,2/3,5), DMS were imaged at 60× magnification on an Olympus Fluoview confocal laser-scanning microscope. Maximum projections of three consecutive optical sections (corresponding to 1 µm total depth) were generated from the original z-stack. Analyses were performed blindly as to genotype. The Puncta Analyzer plugin (written by Barry Wark, modified by Chaichontat (Richard) Sriworarat, and available upon request from Cagla Eroglu at for either ImageJ (NIH; http://imagej.nih.gov/ij/) or FIJI (https://imagej.net/Fiji/Download) was used to count the number of co-localized puncta. This quantification method is based on the fact that pre- and post-synaptic proteins (such as VGluT1 and PSD95) are not within the same cellular compartments of neurons (axons versus dendrites, respectively) and would only appear to partially co-localized at synaptic junctions due to their proximity. This quantification method yields an accurate estimation of the number of synapses both *in vitro* and *in vivo* because it measures co-localization as opposed to staining of a single pre- or postsynaptic protein that often accumulates in extrasynaptic regions during the course of their life cycle. In agreement, numerous previous studies by ourselves and others have shown that synaptic changes observed by this quantification method are verified by techniques such as electron microscopy and electrophysiology ^63, 113, 114–118^. Details of the quantification method have been described previously ^77^. Briefly, 1 µm thick maximum projections are separated into red and green channels, backgrounds are subtracted (rolling ball radius = 50), and thresholds are determined to detect discrete puncta without introducing noise. The minimum pixel size of puncta was set as 4 to remove any background noise. The Puncta Analyzer plugin then uses an algorithm to detect the number of puncta that are in close proximity across the two channels, yielding quantified co-localized puncta.

### Adeno-Associated Virus (AAV) Production

AAV-EF1α-WGA-Dre and AAV-EF1α-(N)Cre-Rox-Stop-Rox-(C)Cre, were produced and purified as described before ^78^. More in detail, AAVs were produced by co-transfecting each AAV vector (15 µg) to the HEK cells with a helper (pAD-delta F6, 30 µg) and the capsid plasmids (15 µg). A total of five 15 cm tissue culture dishes (12 × 10^6^ HEK cells per dish) were used to produce one type of virus. Three days post-transfection, HEK cells were lysed using the cell lysis buffer (Cell lysis buffer: Add 3 ml of 5 M NaCl and 5 ml of 1 M Tris-HCl (pH 8.5) to 80 ml of dH2O. Adjust the pH to 8.5 with NaOH and adjust the volume to 100 ml with dH_2_O. Sterilize by passing through a 0.22-μm filter and store at 4 °C). Then the AAV particles were released from the cells by three cycles of freeze/thaw between dry ice-ethanol and 37 °C water bath and then treated with Benzonase (Novagen, 70664) at 37°C for 30 minutes. Then AAVs were purified using Iodixanol Gradient Ultracentrifugation as described by Addgene (https://www.addgene.org/protocols/aav-purification-iodixanol-gradient-ultracentrifugation/). Briefly, the cell lysates were loaded to the gradients of iodixanol (Sigma D1556, 15%, 25%, 40%, and 60%) in the OptiSeal tubes (Beckman Coulter, Indianapolis, IN). Then the lysates were spin down at 67,000 rpm for 1 hour at 18 °C with the ultracentrifuge (Beckman). The AAV particles were collected by collecting the interface between 40 and 60% iodixanol with a syringe. The collected AAV-containing solution was mixed with ice-cold DPBS and concentrated using the Vivaspin column (100 MWCO) at 4°C. Collected virus particles were aliquoted and stored at -80 °C until use.

### Surgery procedure

Mice were anesthetized with 1.5-2.0% isoflurane mixed with 0.60 L/min of oxygen for surgical procedures and placed into a stereotactic frame (David Kopf Instruments, Tujunga, CA). Meloxicam (2 mg/kg) and topical bupivacaine (0.20 mL) were administered prior to incision. α2δ-1(f/f) and α2δ-1(+/+) mice were used for the behavioral and electrophysiological test as well as for anatomical and synaptic count studies. For these purposes, 50 nL of AAV-EF1α-WGA- Dre were injected bilaterally in the dorsomedial striatum (AP: +0.5 relative to bregma, ML: 1.4 relative to bregma, DV: 2.0 relative to brain surface) and 100 nL of AAV-EF1α-(N)Cre-Rox-Stop- Rox-(C)Cre were injected bilaterally in the frontal and caudal part of the Anterior Cingulate Cortex (AP: +1.5 relative to bregma, ML: 0.12 relative to bregma, DV: 1.5 relative to brain surface; AP: -0.2 relative to bregma, ML: 0.12 relative to bregma, DV: 1.0 relative to brain surface) using a microinjector (Nanoject 3000, Drummond Scientific) at a rate of 1 nL/s.

For optogenetic stimulation experiments, 200 nL of AAV-EF1α-DIO-hChR2(E123T/T15 9C)- eYFP (UNC viral vector core) or AAV-hSyn-DIO-BLINK2-eGFP (Duke viral vector core) were bilaterally injected into the ACC (AP: +0.9 relative to bregma, ML: 0.12 relative to bregma, DV: 1.1 relative to brain surface) and 200 nL of AAV(RETRO2)-EF1α-Cre-WPRE (Duke viral vector core) were bilaterally injected into the DMS (AP: +0.5 relative to bregma, ML: 1.4 relative to bregma, DV: 2.0 relative to brain surface) of WT, α2δ-1(f/f) or α2δ-1(+/+) mice. Custom-made optic fibers (2 - 3 mm length below ferrule, >75% transmittance, 105 μm core diameter) were then implanted directly above the ACC at an angle (AP: +0.5 with respect to bregma, ML: 1.1 with respect to bregma, DV: 1.3 from the brain surface; 25° angle). Fibers were secured in place with dental acrylic adhered to skull screws. Mice were allowed to recover for three weeks after surgery before experimentation.

### Optogenetic Stimulation

For optogenetic stimulations, WT, α2δ-1(f/f), and α2δ-1(+/+) mice expressing ChR2 and implanted with optic fibers were trained for the LP task as described above. After the 3days FR1, mice were trained with the optic fibers attached to the laser source to allow habituation. The training was conducted as described above for the 9 days and in addition, we performed 2 more days with an FR20 schedule. On day 12 the laser was turned on at a frequency of 20 Hz and a power of 7 mW to allow light stimulation for 5 min soon after 5 min without light stimulation. This cycle was repeated 3 times for a total of 30 min. For the PR experiments, mice were trained up to FR10 and on the next day went through either a PR test with optogenetic stimulation or without. After retraining to FR10, the same mice were then tested again on PR with the opposite optogenetic condition.

### Circuit tracing and fiber implant control

The circuit tracing and surgery check were conducted as follow. After the behavioral tests were concluded, mice were anesthetized with 200 mg/kg tribromoethanol (avertin) and euthanized by perfusing with a solution made of TBS with heparin (0.1128g Heparin ammonium salt from porcine intestinal mucosa [Sigma; H6279]) and then with 4% Paraformaldehyde (PFA). Mouse brains were then kept in 4% PFA o.n. at 4°C. The day after brains were rinsed 3 times with TBS, immersed in 30% Sucrose in TBS, and stored at 4°C until they were not floating anymore in the solution. At this time brains were included in a mixture of 30% Sucrose in TBS and Tissue Tek O.C.T. compound (frozen tissue matrix) at a 1:2 ratio and stored at -80°C. Brains were cut into 20 or 50 um coronal sections and stored in a 1:1 mixture of TBS/glycerol at -20 C. Sections were washed in 1x TBS containing 0.2% Triton-X100 (TBST) and blocked in 10% NGS diluted in TBST. For the mice expressing ChR2, BLINK2 or their control with only fiber implants, sections were incubated o.n. with a primary antibody against GFP (1:1000; Millipore, AB16901). In addition, brain sections from α2δ-1(f/f) and α2δ-1(+/+) mice, were also incubated with a primary antibody against RFP (1:2000; Rockland, 600-401-379). Secondary Alexa-fluorophore (488, 594) conjugated antibodies (Invitrogen) were added (1:200 in TBST with 5% NGS) for 2 h at room temperature. Slides were mounted in Vectashield with DAPI (Vector Laboratories, CA) and images were acquired on an Olympus Fluoview confocal microscope using 20X objective at 1.3X Zoom. Mice were excluded if fiber placement was not located in the target site.

### Whole-cell patch-clamp recording

For whole-cell patch-clamp recordings, 4 trained and 4 untrained mice were used per group (WT or α2δ-1(f/f)) to measure miniature excitatory postsynaptic current (mEPSC) before or after training. During all recordings, brain slices were continuously perfused with standard aCSF at RT (∼25°C) and visualized by an upright microscope (BX61WI, Olympus) through a 40x water-immersion objective equipped with infrared-differential interference contrast optics in combination with digital camera (ODA-IR2000WCTRL). Patch-clamp recordings were performed by using an EPC 10 patch-clamp amplifier, controlled by Patchmaster Software (HEKA). Data were acquired at a sampling rate of 50 kHz and low-pass filtered at 6 kHz.

To prepare acute brain slices, after decapitation, the brains were immersed in ice-cold artificial cerebrospinal fluid (aCSF, in mM): 125 NaCl, 2.5 KCl, 3 mM MgCl_2_, 0.1 mM CaCl_2_, 10 glucose, 25 NaHCO_3_, 1.25 NaHPO_4_, 0.4 L-ascorbic acid, and 2 Na-pyruvate, pH 7.3-7.4 (310 mOsmol). Coronal slices containing the ACC were obtained using a vibrating tissue slicer (Leica VT1200; Leica Biosystems). Slices were immediately transferred to standard aCSF (37°C, continuously bubbled with 95% O_2_ – 5% CO2) containing the same as the low-calcium aCSF but with 1 mM MgCl_2_ and 1-2 mM CaCl_2_. After 30 min incubation, slices were transferred to a recording chamber with the same extracellular buffer at room temperature (RT: ∼25°C).

To measure mEPSC, the internal solution contained the following (in mM): 125 K-gluconate, 10 NaCl, 10 HEPES, 0.2 EGTA, 4.5 MgATP, 0.3 NaGTP, and 10 Na-phosphocreatine, pH adjusted to 7.2 – 7.4 with KOH and osmolality set to ∼ 300 mosM. mEPSCs were measured in the aCSF bath solution containing 1 µM tetrodotoxin and 50 µM Picrotoxin at -70 mV in voltage-clamp mode. mEPSCs recorded at -70 mV were detected using Minhee Analysis software (https://github.com/parkgilbong/Minhee_Analysis_Pack). To analyze the frequency, events were counted over 5 minutes of recording. To obtain the average events for each cell, at least 100 non-overlapping events were detected and averaged. The peak amplitude of the average mEPSC was measured relative to the baseline current.

For the data collected only after training, 8 animals were used to measure excitability and 7 animals were used to measure miniature excitatory postsynaptic current (mEPSC). Viral injection to label ACC_->DMS_ neurons was performed as described previously and the animals were sacrificed 6 weeks after the injection. The brain was removed quickly and placed in ice-cold solution bubbled with 95% O2-5% CO2 containing the following (in mM): 194 sucrose, 30 NaCl, 2.5 KCl, 1 MgCl2, 26 NaHCO3, 1.2 NaH2PO4, and 10 D-glucose. After 5 minutes, 250 μm coronal slices were cut with a vibratome (PELCO) and then placed in 35.5°C oxygenated artificial cerebrospinal fluid (aCSF) solution containing the following (in mM): 124 NaCl, 2.5 KCl, 2 CaCl2, 1 MgCl2, 26 NaHCO3, 1.2 NaH2PO4, and 10 D-glucose, pH adjusted to 7.4 with HCl and osmolality set to ∼310 mosM. After 30 minutes, the slices were maintained in aCSF at ∼22 - 23°C for at least 30 min before recording. Following recovery, all recordings were conducted under continuous perfusion of aCSF at 29-30°C, and the pipettès impedances were between 3.5 and 5 MΩ. All recordings were performed with MultiClamp 700B amplifier (Molecular Device) and filtered at 10 kHz and digitized at 20 kHz with a Digidata 1440A digitizer (Molecular Devices).

To measure the excitability, the internal solution contained (in mM) 150 potassium gluconate, 2 MgCl2, 1.1 EGTA, 10 HEPES, 3 sodium ATP, and 0.2 sodium GTP, with pH adjusted to 7.2 with KOH and osmolarity set to ∼300 mosM. Pipettes impedances were between 3.5 and 5 MΩ. The excitability was measured in current-clamp mode by injection of current between -300 and 400 pA. Each step was 100LJpA with a duration of 1 s. The number of spikes for each depolarizing step was counted by peak detection software in pCLAMP10 (Molecular Devices).

To measure miniature excitatory postsynaptic current (mEPSC), the internal solution contained the following (in mM): 120 cesium methanesulfonate, 5 NaCl, 10 tetraethylammonium chloride, 10 HEPES, 4 lidocaine N-ethyl bromide, 1.1 EGTA, 4 magnesium ATP, and 0.3 sodium GTP, pH adjusted to 7.2 with CsOH and osmolality set to ∼ 300 mosM. mEPSCs were measured in the aCSF bath solution containing 1 µM tetrodotoxin and 50 µM Picrotoxin at -70 mV in voltage-clamp mode. The amplitudes of mEPSCs over -10 pA by the peak detection software in pCLAMP10 were counted.

### Quantification and statistical analysis

All statistical analyses were performed in GraphPad Prism 8. Sample size and specific statistical tests for each experiment are indicated in the figure legend for each experiment. Exact adjusted p-values are listed in the figures for each experiment, where absent the difference was not significant (p>0.05). the details of the statistical analysis are reported in figure legends. The significance for all the quantifications is *p<0.05, **p<0.01, ***p<0.001, and ****p<0.0001. The RNA sequencing data were analyzed using the Benjamini-Hochberg correction and the false discovery rate (FDR) was utilized to evaluate differences among data sets. Where indicated, the Unpaired two-tailed t-test were run using Welch’s correction and then a correction for multiple comparison was applied using Hom-Sidak method with an alpha threshold of 0.05 for adjusted p value. A Geisser-Greenhouse correction was used for both One-way and Two-way ANOVA analyses. Sample sizes were determined based on previous experience for each experiment to yield high power to detect specific effects. No statistical methods were used to predetermine sample size.

## Data availability

Further information and requests for resources and reagents can be directed to the Lead Contacts, Francesco Paolo Ulloa Severino, Henry Yin, and Cagla Eroglu. The reagents, data, and code generated in this study are available without restriction.

---

**Figure Supplementary 1.**
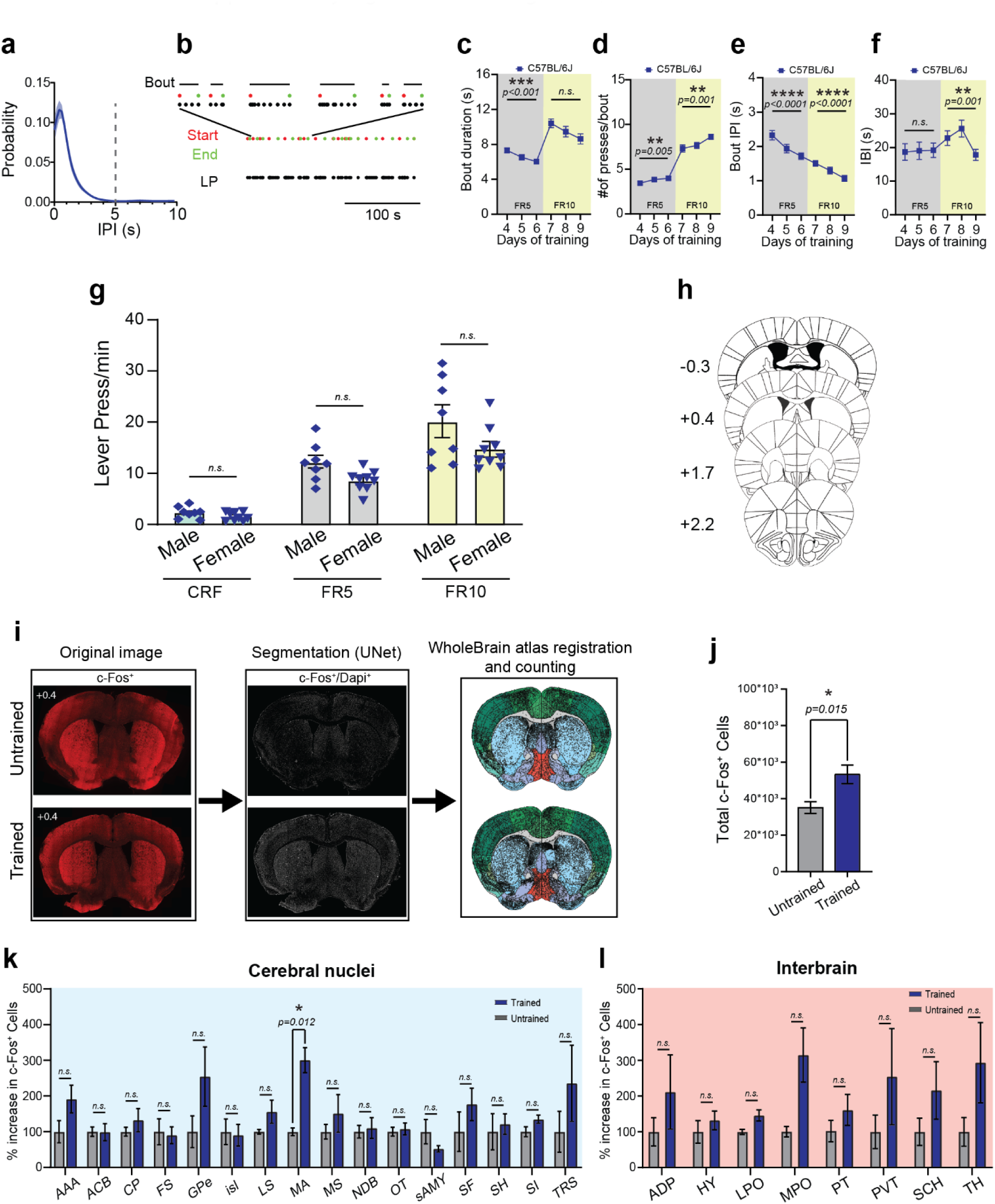
**a**, Probability of inter-press interval (IPI) utilized to identify lever press bouts. **b**, Example of identified Start and End of the lever press bouts within a behavioral session. **c**, Bout Duration (*RM one-way ANOVA*. FR5 *[F (1.82, 29.1) = 16.3], p < 0.0001*; FR10 *[F (1.32, 21.0) = 5.71], p = 0.019*). *Holm-Sidak’s multiple comparisons adjusted p-value reported in figure*. **d**, Number of presses per bout (*RM one-way ANOVA*. FR5 *[F (1.98, 31.6) = 7.86], p = 0.001*; FR10 *[F (1.96, 31.3) = 11.7], p < 0.001*). *Holm-Sidak’s multiple comparisons adjusted p-value reported in figure*. **e**, Bout IPI (*RM one-way ANOVA.* FR5 *[F (1.73, 27.7) = 36.0], p < 0.0001*; FR10 *[F (1.78, 28.4) = 21.8], p < 0.0001*). *Holm-Sidak’s multiple comparisons adjusted p-value reported in figure*. **f**, Inter Bout Interval (*RM one-way ANOVA [F (2.32, 37.1) = 3.22], p=0.045*). *Holm-Sidak’s multiple comparisons adjusted p-value reported in figure*. **g**, Quantification of lever press/min in male (n = 8) versus female (n = 9) mice. *RM Two-way ANOVA, main effect of ratio [F (1.142, 17.13) = 112.3, p<0.0001]* and no effect of sex *[F (1,15) = 4.155, p=0.059]* nor interaction *[F (2,30) = 2.442, p=0.104]*. **h**, Stereotaxic coordinates of the brain sections used for the c-Fos analysis. **i**, Image processing flow chart with examples for a trained and an untrained animal. **j**, Total numbers of c-Fos^+^ cells in Untrained (35.1 ± 3.2 *10^3^ c-Fos+ cells) and Trained mice (53.2 ± 5.1 *10^3^ c-Fos+ cells). n = 6 mice per condition; mean of 4 sections per mouse. *Unpaired two-tailed Mann-Whitney test [U = 3]*. **k**, Bar plot of c-Fos^+^ cells for the cerebral nuclei regions. *Multiple unpaired t-test with Welch correction. Multiple comparisons using Holm-Sidak method; alpha = 0.05 for adjusted p-value*. MA (300.54 ± 35.11%, *[t (6.26) = 6.0]*). n = 4-6 mice per condition. **l**, Bar plot of c-Fos^+^ cells for the Interbrain regions. *Multiple unpaired t-test with Welch correction. Multiple comparisons using Holm-Sidak method; alpha = 0.05 for adjusted p-value*. n = 4-6 mice per condition. For all graphs: Data shown as mean ± s.e.m. alpha = 0.05.

**Figure Supplementary 2.**
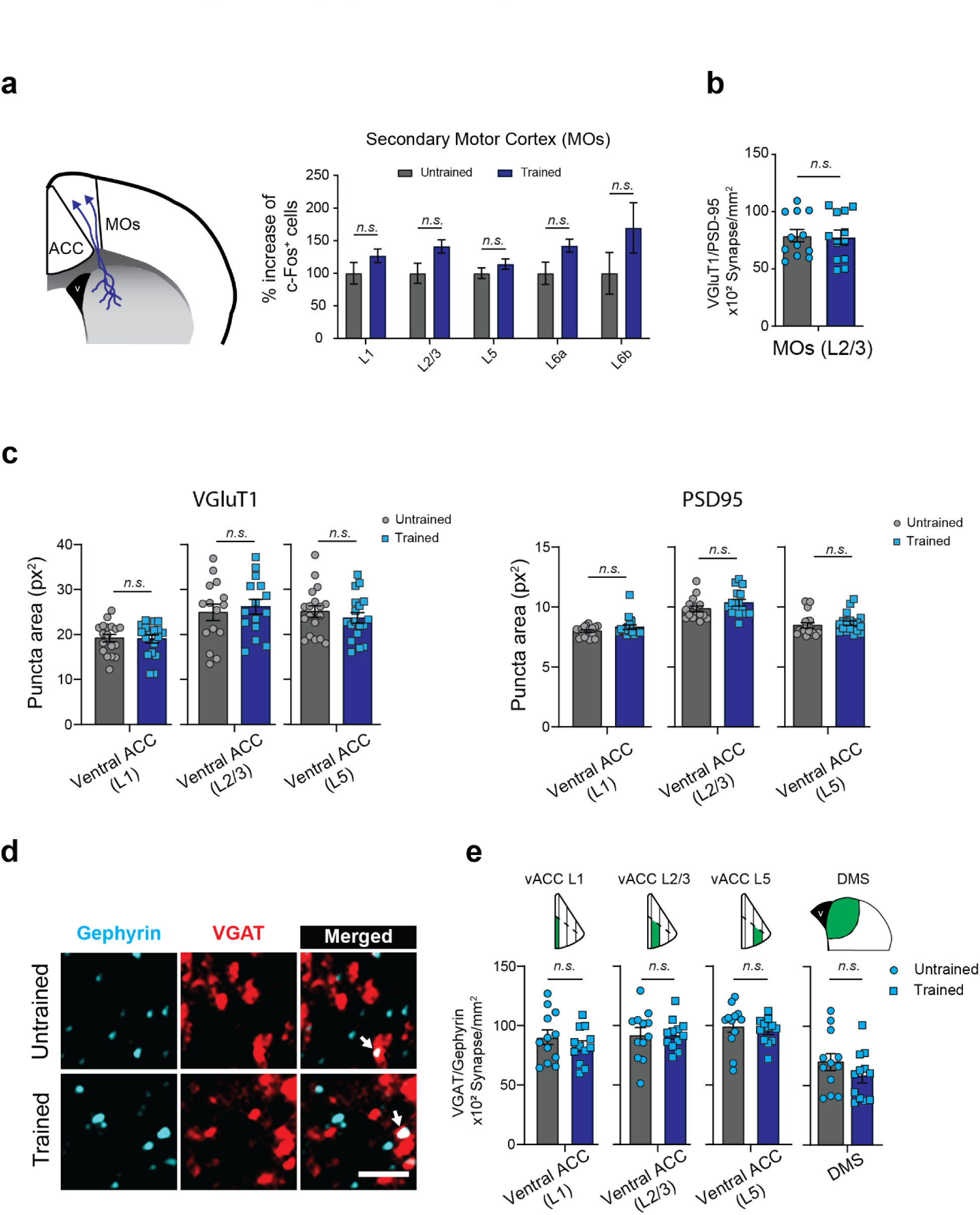
**a**, *Left.* Schematic representation of L2/3 neurons projecting from the MOs to the DMS. *Right.* Layer-specific count of c-Fos^+^ cells in the MOs. *Multiple unpaired t-test with Welch correction. Multiple comparisons using Holm-Sidak method; alpha = 0.05 for adjusted p-value*. **b**, Quantification of VGlut1/PSD95 co-localized puncta in layer 2/3 of MOs (n = 4 mice per condition, 3 images per mouse). *Unpaired Two-tailed t-test with Welch’s correction [t (22) = 0.17].* **c**, Quantification of synaptic puncta area for VGluT1 and PSD95 in Untrained and Trained mice. *Unpaired Two-tailed t-test*. L1 Untrained VGluT1 (19 ± 0.83), Trained VGluT1 (19 ± 0.88), *[t (34) = 0.17]*; PSD95 Untrained (8.0 ± 0.1), Trained (8.3 ± 0.19), *[t (34) = 1.5]*; L2/3 Untrained VGluT1 (25 ± 1.8), Trained VGluT1 (26 ± 1.6), *[t (28) = 0.5]*; PSD95 Untrained (9.8 ± 0.24), Trained (10 ± 0.29), *[t (28) = 1.4]*; L5 Untrained VGluT1 (25 ± 1.3), Trained VGluT1 (24 ± 1.2), *[t (34) = 0.84]*; PSD95 Untrained (8.5 ± 0.22), Trained (8.6 ± 0.19), *[t (34) = 0.6]*. **d**, Representative images for Untrained and Trained mice of Gephyrin and VGAT staining. The *arrows* in the merged channel indicate co-localized puncta. *Scale bar 20 µm*. **e**, Quantification of Gephyrin/VGAT co- localized puncta in ACC (L1, L2/3 and L5). *Unpaired Two-tailed t-test.* L1 Untrained (90 ± 6.1), Trained (83 ± 4.3) *[t (22) = 0.99]*; L2/3 Untrained (92 ± 6.0), Trained (92 ± 3.7) *[t (22) = 0.00028]*; L5 Untrained (100 ± 5.4), Trained (96 ± 3.2) [t (22) = 0.67]; and DMS (Untrained (70 ± 7.1), Trained (58 ± 5.8), *[t (22) = 1.3].* n = 4 mice per condition, 3 images per mouse. For all graphs: Data shown as mean ± s.e.m. alpha = 0.05.

**Figure Supplementary 3.**
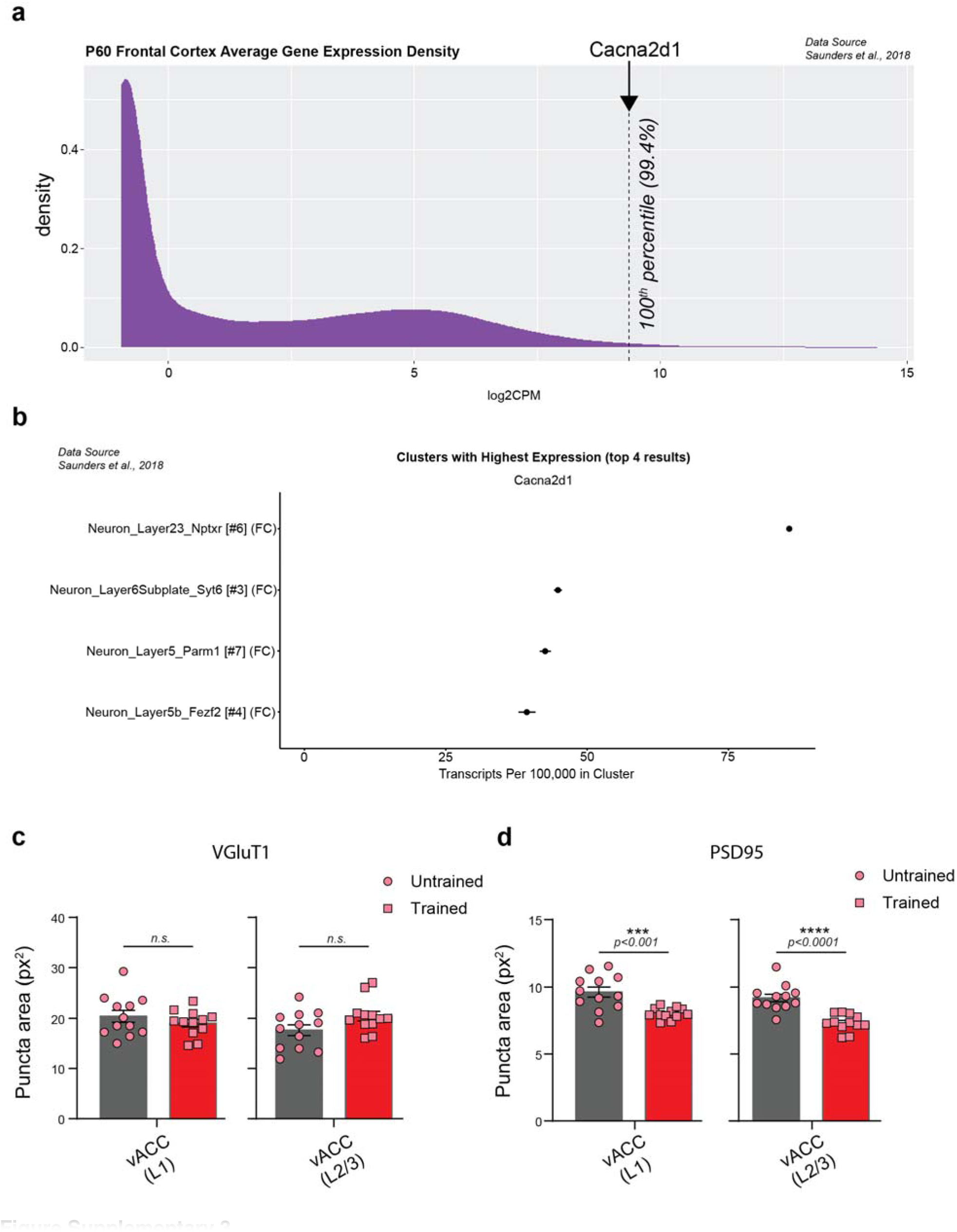
**a**, Expression density (log_2_CPM) of Cacna2d1 gene (encoding for α2δ-1) in the prefrontal cortex of adult mice (P60). Data mined from Saunders et al., 2018. **b**, Cacna2d1 expression level across layers of the prefrontal cortex. Data mined from Saunders et al., 2018. **c**, Quantification of synaptic puncta area for VGluT1 in Untrained and Trained α2δ-1 KO mice. *Unpaired Two-tailed t-test.* L1 Untrained VGluT1 (20 ± 1.2), Trained VGluT1 (19 ± 0.75), *[t (22) = 0.98]*; L2/3 Untrained VGluT1 (18 ± 1.1), Trained VGluT1 (20 ± 0.95), *[t (22) = 1.9]*. **d**, Quantification of synaptic puncta area for PSD95 in Untrained and Trained α2δ-1 KO mice. *Unpaired Two-tailed t-test.* L1 Untrained PSD95 (9.6 ± 0.37), Trained PSD95 (8.0 ± 0.12), *[t (22) = 4.1]*; L2/3 Untrained PSD95 (9.2 ± 0.28), Trained PSD95 (7.3 ± 0.18), *[t (22) = 5.5];*

**Figure Supplementary 4.**
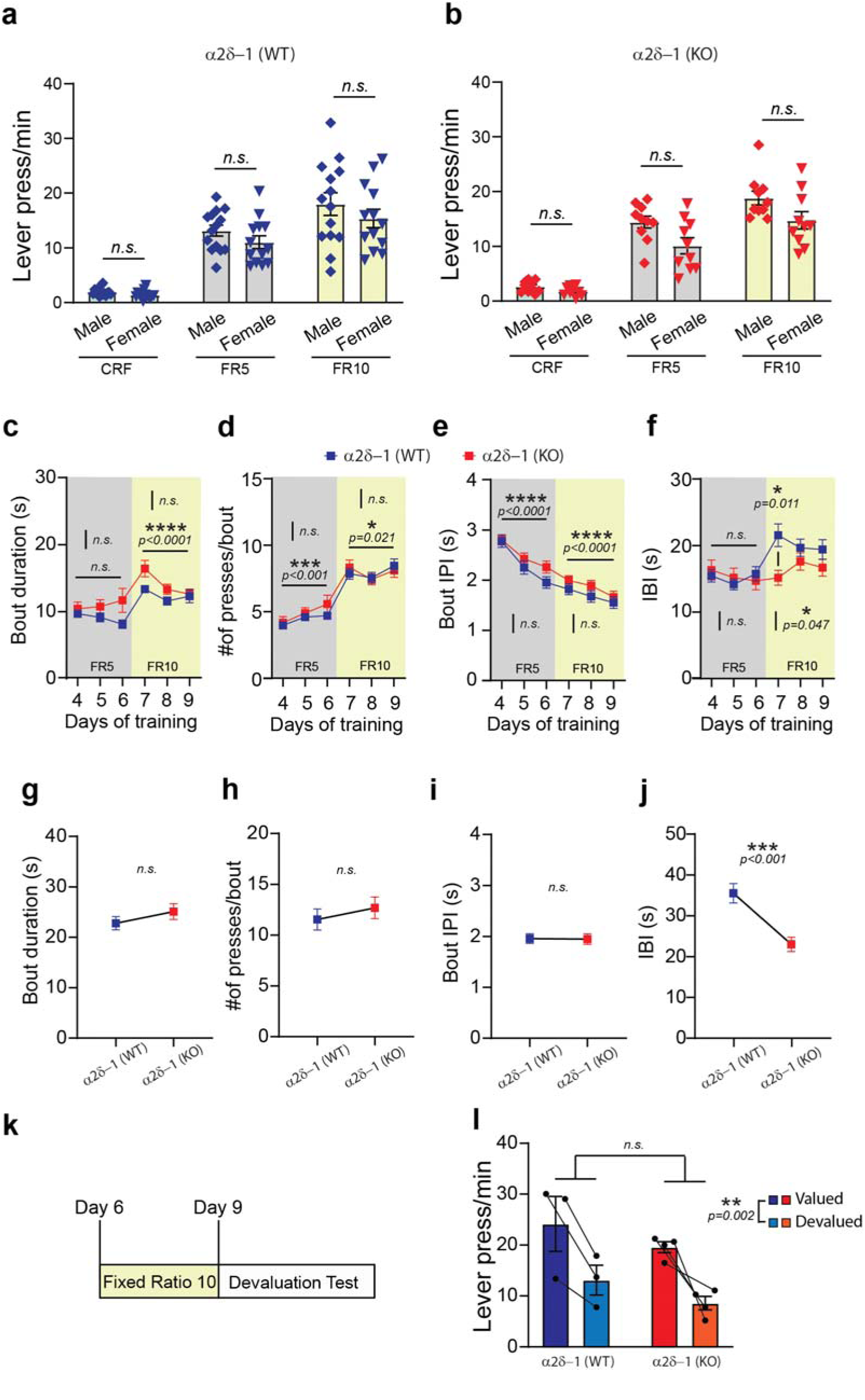
**a**, Quantification of lever press/min in α2δ-1 WT male (n = 14) versus female (n = 13) mice. α2δ-1 WT: RM Two-way ANOVA, main effect of ratio [F (1.31, 32.75) = 126.6, p < 0.0001] and no effect of sex [F (1, 25) = 1.6, p = 0.217] nor interaction [F (2, 50) = 0.6066, p = 0.549]. **b**, Quantification of lever press/min in α2δ-1 KO male (n = 10) versus female (n = 10) mice. α2δ-1 KO: RM Two-way ANOVA, main effect of ratio [F (1.707, 30.73) = 191.4, p < 0.0001], main effect of sex [F (1, 18) = 5.317, p = 0.033] and interaction [F (2, 36) = 3.285, p = 0.048]. Multiple comparisons using Holm-Sidak method; alpha = 0.05 for adjusted p-value showed no sexual dimorphism, CRF [t (17.26) = 1.97, p = 0.18], FR5 [t (16.68) = 2.37, p = 0.088], FR10 [t (17.16) = 1.99, p = 0.174]. **c**, Bout Duration (RM Two-way ANOVA. <u>FR5</u>: No effect of Days [F (1.308, 58.87) = 0.03069, p = 0.915] nor Genotype [F (1, 45) = 3.407, p = 0.071] and no interaction [F (2, 90) = 2.435, p = 0.093]). <u>FR10:</u> Main effect of Days [F (1.778, 79.99) = 15.93], no effect of Genotype [F (1, 45) = 2.796, p = 0.101] and interaction [F (2, 90) = 3.810, p = 0.026], Multiple comparison showed no effect between genotypes). **d**, Number of presses per bout (RM Two-way ANOVA. <u>FR5</u>: Main effect of Days [F (1.324, 59.57) = 10.18] no effect of Genotype [F (1, 45) = 1.026, p = 0.316] and no interaction [F (2, 90) = 1.181, p = 0.312]). <u>FR10:</u> Main effect of Days [F (1.729, 77.82) = 4.322] no effect of Genotype [F (1, 45) = 8.166e-005, p = 0.993] and no interaction [F (2, 90) = 1.180, p = 0.312]). **e**, Bout IPI (RM Two-way ANOVA. <u>FR5</u>: Main effect of Days [F (1.559, 70.16) = 116.8] no effect of Genotype [F (1, 45) = 1.031, p = 0.315] and significant interaction [F (2, 90) = 4.791, p = 0.010]. Multiple comparison showed no differences between genotypes). <u>FR10:</u> Main effect of Days [F (1.637, 73.65) = 39.08] no effect of Genotype [F (1, 45) = 1.103, p = 0.299] and no interaction [F (2, 90) = 1.209, p = 0.303]). **f**, Inter Bout Interval (RM Two-way ANOVA. <u>FR5</u>: No effect of Days [F (1.706, 76.76) = 2.279, p = 0.117] no effect of Genotype [F (1, 45) = 0.03223, p = 0.858] and no interaction [F (2, 90) = 2.044, p = 0.135]). FR10: No effect of Days [F (1.488, 66.95) = 0.2467, p = 0.715], main effect of Genotype [F (1, 45) = 4.166, p = 0.047] and significant interaction [F (2, 90) = 3.916, p = 0.023]). **g**, Bout Duration in α2δ-1 WT (23 ± 1.3) and KO (25 ± 1.6), Unpaired Two-tailed t-test. [t (39) = 1.1, p = 0.266]. **h**, Number of presses per bout in α2δ-1 WT (12 ± 1.0) and KO (13 ± 1.1), Unpaired Two-tailed t-test. [t (39) = 0.77, p = 0.448]). **i**, Bout IPI in α2δ-1 WT (2 ± 0.09) and KO (1.9 ± 0.10), Unpaired Two-tailed t-test. [t (39) = 0.079, p = 0.937]. **j**, Inter Bout Interval in α2δ-1 WT (36 ± 2.4) and KO (23 ± 1.8), Unpaired Two-tailed t-test. [t (39) = 4.1, p<0.001]. **k**, Schematic representation of the devaluation test schedule. **l**, Lever press/min in valued and devalued states after pre-feeding. RM Two-way ANOVA. Main effect of value state [F (1, 5) = 35.48], no effect of genotype [F (1, 5) = 1.750, p = 0.243] nor interaction [F (1, 5) = 0.0001262, p = .991]. For all the graphs: Data shown as mean ± s.e.m. Multiple comparisons using Holm-Sidak method; alpha = 0.05 for adjusted p-value

**Figure Supplementary 5.**
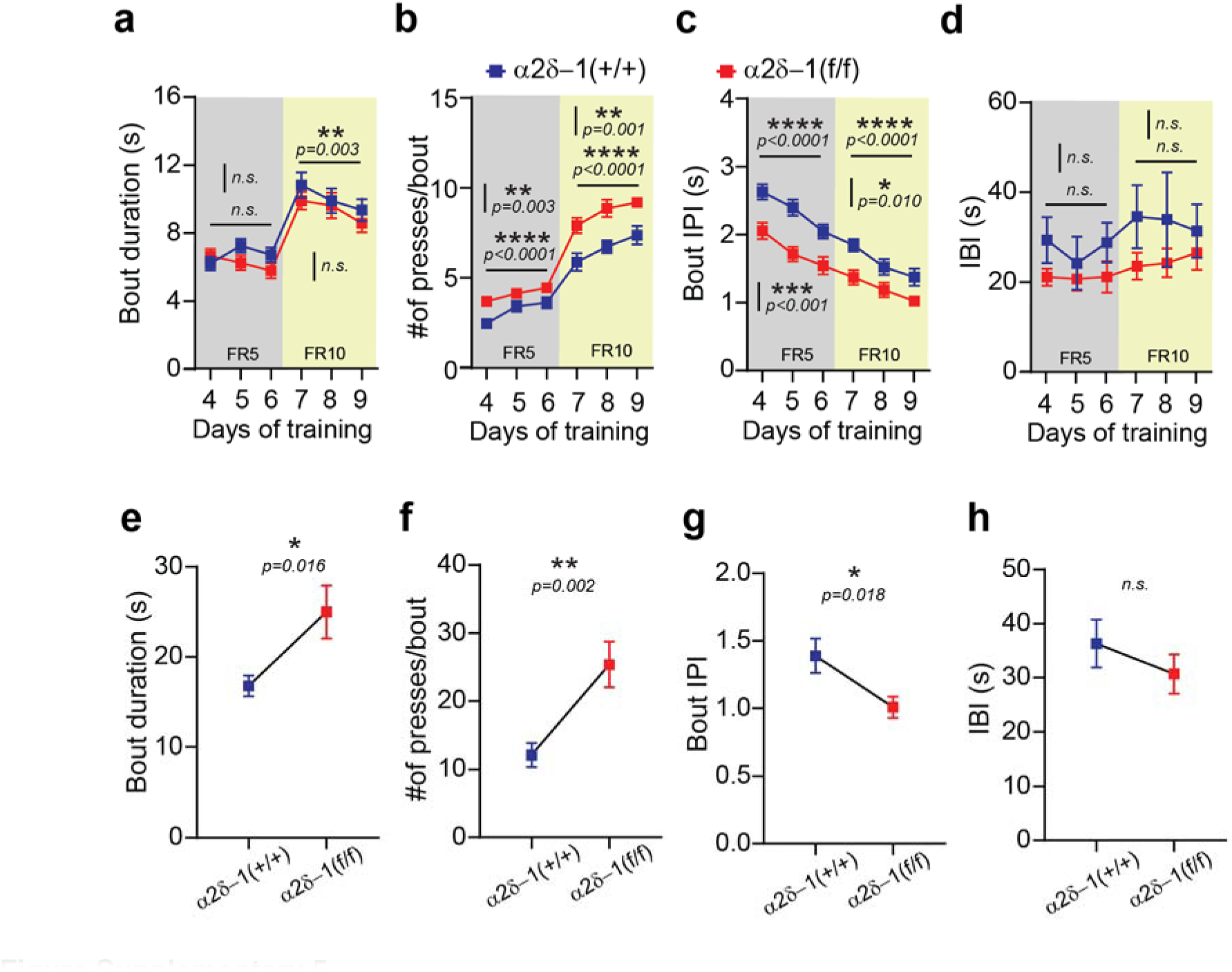
**a**, Bout Duration (RM Two-way ANOVA. <u>FR5:</u> no effect of Days [F (1.348, 29.66) = 1.629, p = 0.215] nor Genotype [F (1, 22) = 0.9778, p = 0.333] and significant interaction [F (2, 44) = 4.105, p = 0.023]). Multiple comparison showed no differences between genotypes. <u>FR10:</u> Main effect of Days [F (1.760, 38.72) = 7.238], no effect of Genotype [F (1, 22) = 0.6237, p = 0.438] and no interaction [F (2, 44) = 0.4177, p = 0.661]). **b**, Number of presses per bout (RM Two-way ANOVA. <u>FR5:</u> Main effect of Days [F (1.809, 39.81) = 21.29] and Genotype [F (1, 22) = 11.04] and no interaction [F (2, 44) = 1.563, p = 0.221]. <u>FR10:</u> Main effect of Days [F (1.969, 43.32) = 19.06] and Genotype [F (1, 22) = 13.08] and no interaction [F (2, 44) = 0.2286, p = 0.796]). **c**, Bout IPI (RM Two-way ANOVA. <u>FR5:</u> Main effect of Days [F (1.543, 33.95) = 34.17] and of Genotype [F (1, 22) = 16.07] and no significant interaction [F (2, 44) = 0.9428, p = 0.397]. <u>FR10:</u> Main effect of Days [F (1.956, 43.04) = 38.95] and Genotype [F (1, 22) = 7.784] and no interaction [F (2, 44) = 1.322, p = 0.277]). **d**, Inter Bout Interval (RM Two-way ANOVA. <u>FR5:</u> No effect of Days [F (1.769, 38.92) = 0.5621, p = 0.553] no effect of Genotype [F (1, 22) = 1.810, p = 0.192] and no interaction [F (2, 44) = 0.4030, p = 0.671]. <u>FR10:</u> No effect of Days [F (1.804, 39.70) = 0.001295, p = 0.997], no effect of Genotype [F (1, 22) = 1.130, p = 0.299] and no interaction [F (2, 44) = 0.5863, p = 0.561]). **e**, Bout Duration in α2δ-1 (+/+) (17 ± 1.2) and α2δ-1 (f/f) (25 ± 2.9), Unpaired Two-tailed t-test. [t (22) = 2.6,]. **f**, Number of presses per bout in α2δ-1 (+/+) (12 ± 1.7) and α2δ-1 (f/f) (25 ± 3.4), Unpaired Two-tailed t-test. [t (22) = 3.5]. **g**, Bout IPI in α2δ-1 (+/+) (1.4 ± 0.13) and α2δ-1 (f/f) (1.0 ± 0.08), Unpaired Two-tailed t-test. [t (22) = 2.5]. **h**, Inter Bout Interval in α2δ-1 (+/+) (36 ± 4.4) and α2δ-1 (f/f) (31 ± 3.6), Unpaired Two-tailed t-test. [t (22) = 0.99, p = 0.335]. For all the graphs: Data shown as mean ± s.e.m. Multiple comparisons using Holm-Sidak method; alpha = 0.05 for adjusted p-value

**Figure Supplementary 6.**
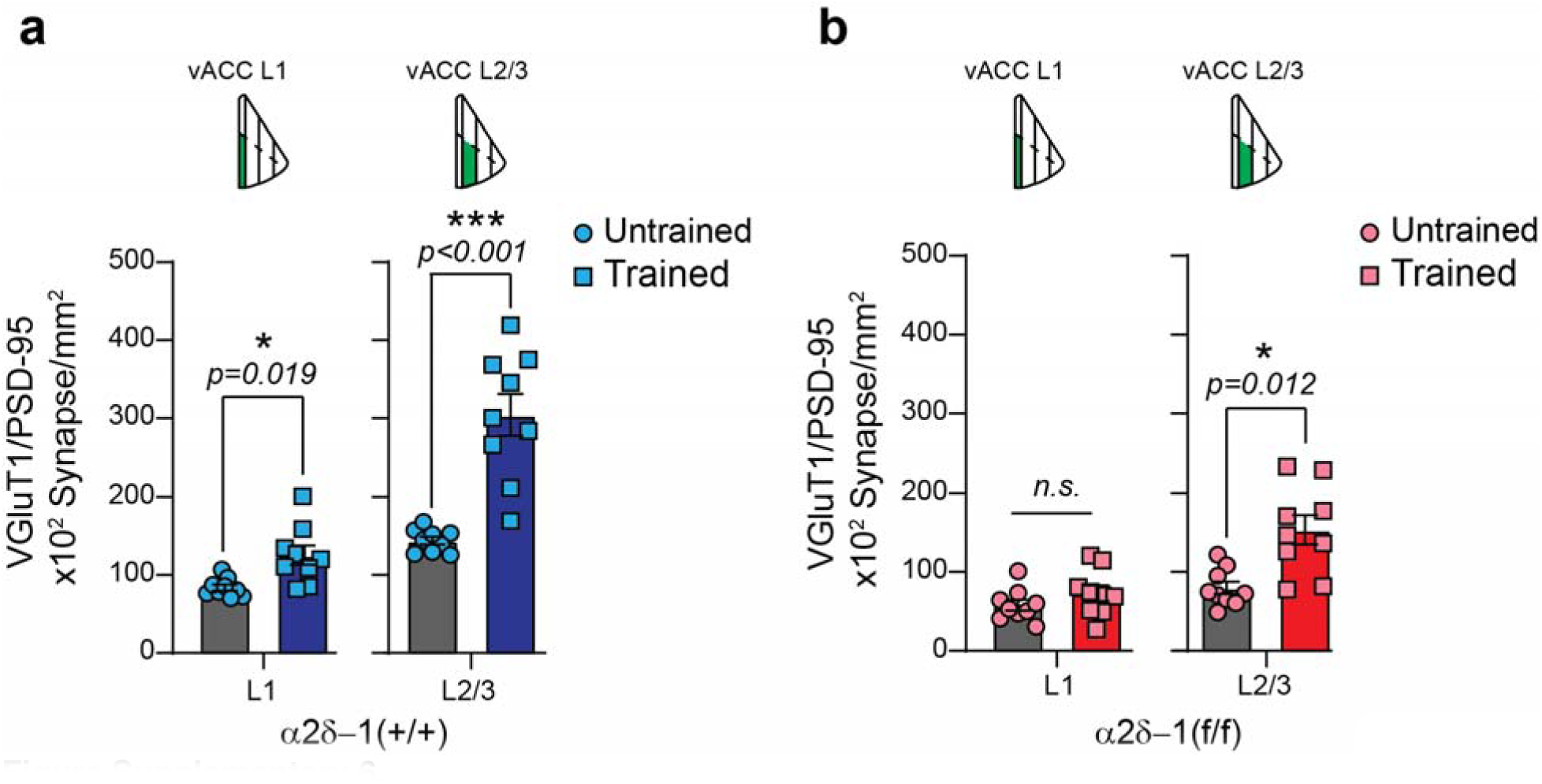
**a**, Quantification of VGlut1/PSD95 co-localized puncta in L1 and L2/3 of vACC for untrained and trained α2δ-1 (+/+) (n = 3 mice per condition, 3 images per mouse). *Multiple unpaired Two-tailed t-test with Welch’s correction, multiple comparison with Holm-Sidak method alpha = 0.05 for adjusted p-value.* L1 *[t (9.6) = 3.2];* L2/3 *[t (8.5) = 5.8].* **b**, Quantification of VGlut1/PSD95 co-localized puncta in L1 and L2/3 of vACC for untrained and trained α2δ-1 (f/f) (n = 3 mice per condition, 3 images per mouse). *Multiple unpaired Two-tailed t-test with Welch’s correction, multiple comparison with Holm-Sidak method alpha = 0.05 for adjusted p-value.* L1 *[t (14.14) = 1.2, p = 0.408];* L2/3 *[t (10.79) = 3.6].* For all the graphs: Data shown as mean ± s.e.m.

**Figure Supplementary 7.**
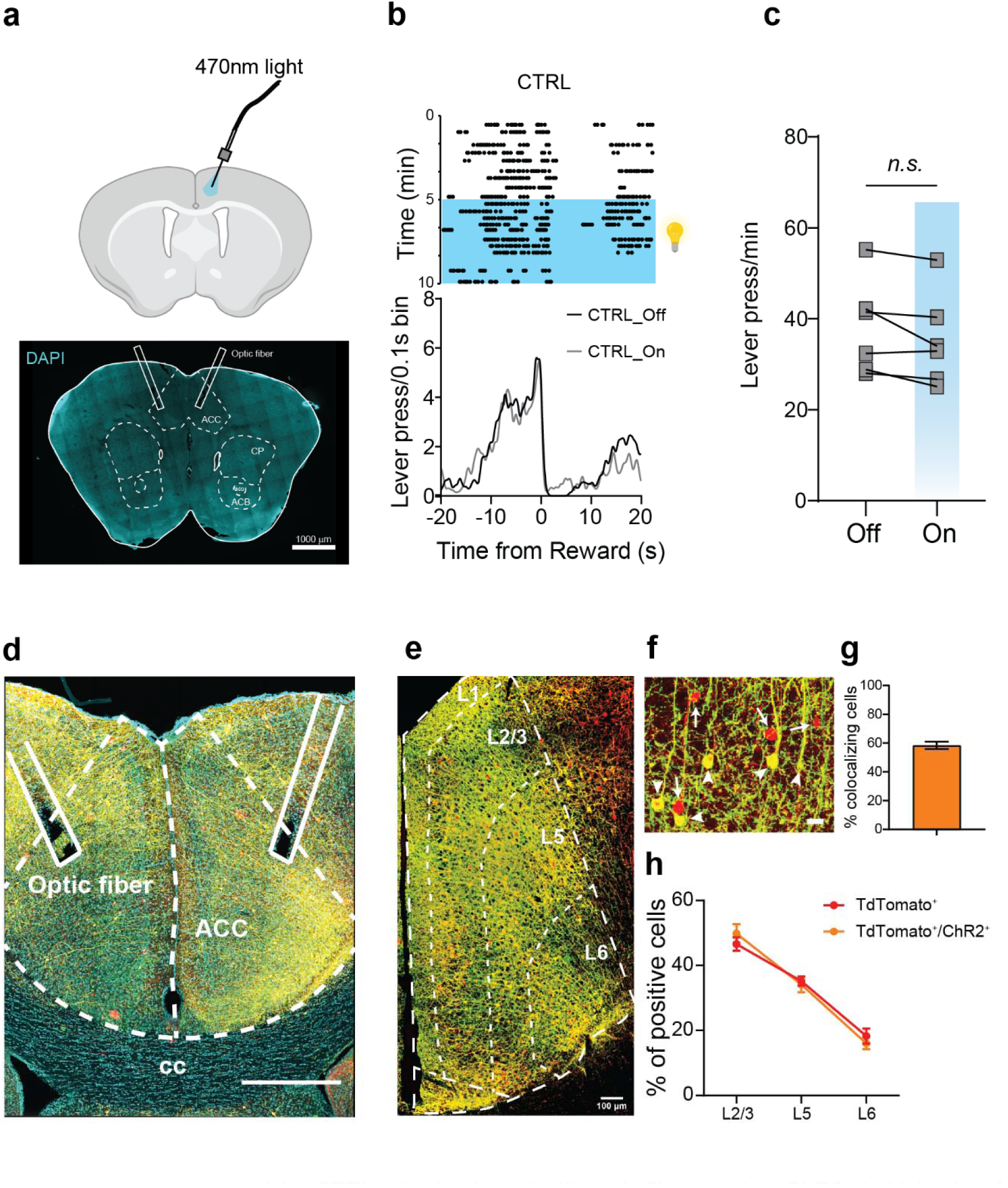
**a**, Schematic representation of the CTRL mice implanted with optic fibers and no ChR2 viral injection. **b**, Example peri-reward raster histogram during time of light-Off and light-On. **c**, Lever press/min for CTRL mice (n = 6) during light-Off (38 ± 4.2) and light-On (35 ± 4.2). *Paired Two-tailed t-test [t (5) = 2.2, p = 0.079]*. **d**, Anatomical confirmation of viral expression and fiber placement in a representative example of an α2δ-1(f/f) mouse expressing the Rox-Cre and the Cre-dependent ChR2 in ACC_->DMS_ neurons. **e**, Representative image of ACC_->DMS_ neurons across layers positive for tdTomato and ChR2. **f**, Magnified image of ACC_->DMS_ neurons labeled by tdTomato (arrow) and ChR2, yellow neurons (arrow heads) are the ones expressing both molecules. **g**, Quantification of colocalized cells within the ACC (h) Distribution of tdTomato^+^ and tdTomato/ChR2^+^ cells across layers. For all graphs: Data shown as mean ± s.e.m. alpha = 0.05.

